# Cooperative colonization of the host and pathogen dissemination involves stochastic and spatially structured expression of virulence traits

**DOI:** 10.1101/2024.03.12.584633

**Authors:** Nieves López-Pagán, José S. Rufián, Julien Luneau, María-Antonia Sánchez-Romero, Laurent Aussel, Simon van Vliet, Javier Ruiz-Albert, Carmen R. Beuzón

## Abstract

Bacteria respond to changing environments by altering gene expression. Some responses display probabilistic cell-to-cell variation within isogenic populations. A few paradigmatic examples in animal pathogens have demonstrated that this phenotypic heterogeneity has biological relevance for virulence.

We investigate single-cell flagellar expression in relation to type III secretion expression in the plant pathogen *Pseudomonas syringae* and describe that both systems undergo phenotypic heterogeneity throughout plant colonization. We establish that high expression of these system carries growth penalties. Stochastic, spatial and time factors shape dynamics of a phenotypically diverse population which displays division of labor during colonization: T3SS^ON^ bacteria effectors act as ‘common goods’ to suppress immunity, allowing the increase of motile bacteria that actively leave the infected tissue before necrosis. This study provides a comprehensive view of how processes underlying bacterial specialization play out in the context of complex and changing environments of biological and applied relevance such as host colonization.

## INTRODUCTION

Flagellar motility is an important trait for colonization processes and environmental adaptation. Mutants in flagellar regulation, biogenesis and/or modification are usually affected in their ability to successfully navigate the environment. Members of the genus *Pseudomonas* can colonize niches as diverse as soil, plant, or animal tissues. The *Pseudomonas syringae* species complex includes most of those pathogenic to plants, causing diseases in a wide range of plant hosts, including many of agricultural relevance^1^. Bacteria from this complex can live on the surface of leaves as epiphytes, but most favor endophytic colonization, using wounds and stomata to enter the leaf apoplast. Flagellar motility confers advantages to epiphytic populations of *P. syringae* pv. syringae^2^, and facilitates active bacterial entry within the leaf for many *P. syringae* pathovars^3–8.Once^ within the apoplast, flagellar motility is not required for systemic spread^3^, while flagellin, the main component of the flagellar pilus, triggers immunity upon recognition by plant response receptors^9^. Fittingly, flagellar expression is downregulated once bacteria enter the apoplast^10–12^

Flagellar biosynthesis is classically depicted as a deterministic program with a complex and tiered regulatory hierarchy, where promoters are sequentially activated maintaining this state throughout active growth. However, single-cell level studies have revealed stochastic activation pulses and bimodal expression patterns in the well-characterized flagellar system of the animal pathogen *Salmonella enterica*, with clonal populations displaying phenotypic heterogeneity in laboratory media and ON and OFF states for the expression of flagellar genes^13–20^. As for *P. syringae*, conservation of gene arrangement and promoter motifs with *P. putida* supports a three-tiered hierarchy of transcriptional regulation of flagellar synthesis^21^, with transcriptional regulator FleQ at the top^22^, however the regulatory cascade remains mostly uncharacterized.

Apoplastic *P. syringae* use a type III secretion system (T3SS) to introduce bacterial effectors into the cytosol of surrounding host cells to suppress plant immunity, including flagellin-triggered defense responses, thus allowing bacterial proliferation and disease development (reviewed in Schreiber et al.^23^). Expression of T3SS genes requires the extra cytoplasmic function (ECF) sigma factor HrpL^24^. Previously, we found that a mutant lacking HrpL (*ΔhrpL*) displays increased swimming motility in minimal apoplast-mimicking medium, while a mutant lacking HrpV, a repressor of *hrpL* expression^25^, displays reduced motility^26^, suggesting a potential counter-regulation between T3SS expression and flagellar motility. We later established that *P. syringae* T3SS genes display phenotypic heterogeneity during plant colonization and stochastic bistable expression with reversible ON and OFF states upon induction during growth in minimal apoplast-mimicking medium^27^. This was a first example of phenotypic heterogeneity for a virulence trait in plant pathogenic bacteria.

Here, we show that flagellar gene expression consistently displays phenotypic heterogeneity in *P. syringae*, particularly during growth within the plant apoplast. While the average expression level of flagellin of the apoplastic population is downregulated, as previously described, a significant percentage of the bacterial population expresses the gene at high levels, particularly during late stages of colonization of this niche. We analyze flagellar expression and T3SS expression simultaneously and identify all potential single-cell ON/OFF combinations, revealing an increasing phenotypic complexity arising amid otherwise clonal bacterial populations. Even so, T3SS^ON^/Flagella^OFF^ and T3SS^OFF^/Flagella^ON^ subpopulations were generally more abundant, suggesting a degree of counter-regulation between these two loci. We show that expression of either of these systems has an impact on bacterial growth, at the population and/or single-cell cell level. Analysis of the spatial and dynamic distribution of expression of these two traits within the apoplastic microcolony through plant colonization show how T3SS^ON^ bacteria are more abundant in the early stages of the infection and close to the host cell surface, with Flagella^ON^ becoming more frequent at later stages and further away from the host cell. These expression patterns display a spatially structured distribution within the host tissue. Finally, we show how following initial bacterial multiplication within the apoplast, Flagella^ON^ bacteria selectively and actively exit the plant leaf, from early in the infection process, prior to the onset of disease symptoms. We propose a division of labor model in which T3SS^ON^ bacteria share effector activity as common goods, complementing T3SS^OFF^ bacteria, while Flagella^ON^ bacteria actively swim out of the tissue prior to tissue collapse.

## RESULTS

### Flagellar expression displays phenotypic heterogeneity in *P. syringae*

We generated a transcriptional fusion to *GFP3* of the flagellin-encoding *fliC* gene within its native chromosome location (preserving genome context and gene function) in the model bean pathogen *P. syringae* pv. phaseolicola 1448A^28^. We used this strain to monitor single-cell flagellin expression in laboratory media using fluorescence confocal microscopy and flow cytometry (**Fig. 1A**). We found that while the *fliC::GFP3* fusion is expressed in the majority of the bacterial cells (Flagella^ON^), fluorescence levels vary considerably cell-to-cell (**Fig. 1A**). Furthermore, a proportion of the population does not display any fluorescence (Flagella^OFF^) and can be visualized only by membrane staining, overlapping with the non-fluorescent control strain in the flow cytometry analysis (**Fig. 1A**). Although phenotypic heterogeneity is common to both both rich (LB) and plant apoplast-mimicking Hrp-inducing (HIM) media (**Fig. 1A**), population-average *fliC* expression levels are lower in LB (**Fig. 1B**).

**Figure 1.**
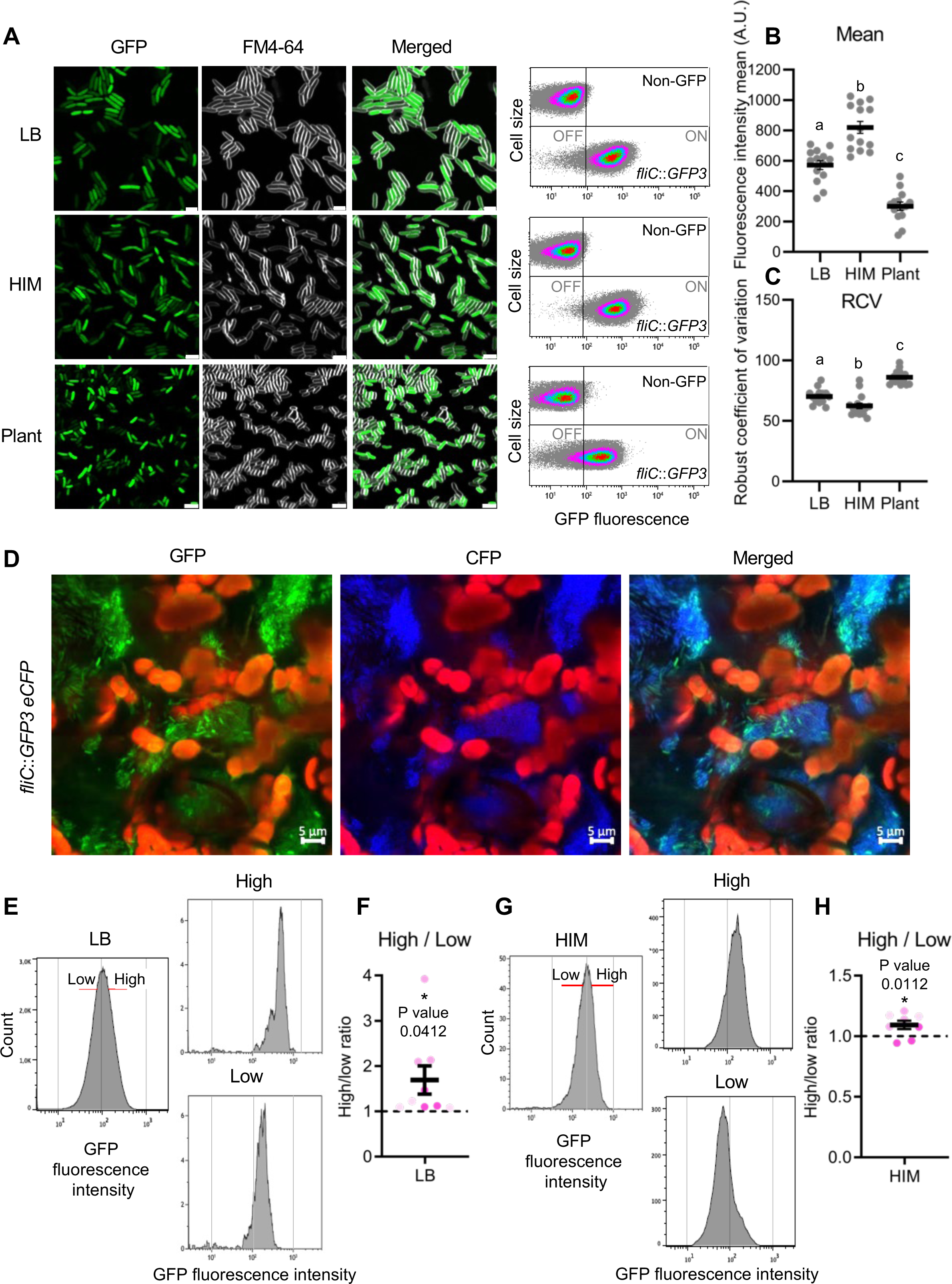
Flagella display phenotypic heterogeneity in *Pseudomonas syringae*. (A) Confocal microscopic images and flow cytometry analysis of a strain carrying a chromosome-located *fliC*::*GFP3* transcriptional fusion either grown overnight in LB (upper panels), for 24 hours in HIM (central panels), or extracted from bean leaf apoplasts 4 days post inoculation (dpi) by infiltration with 5×10^4^ CFU/ml (bottom panels). Microscopy images show in the GFP channel the fluorescence of GFP as reporter of the *fliC* gene expression. FM4-64 channel (shown in greyscale) display fluorescence corresponding to membrane staining and merged panels show the combined GFP and FM4-64 channels. Scale bars correspond to 2 µm. Contrast and brightness were adjusted to improve visualization but were kept constant across the different conditions and channels. Flow-cytometry analysis of the *fliC*:*:GFP3* strain is shown as dot plots representing GFP fluorescence intensity *versus* cell size. Data are represented as arbitrary units in logarithmic scale. Data displayed corresponds to that collected for at least 100,000 events per sample. The non-GFP graph shows autofluorescence levels displayed by the wild type strain not carrying any fluorescent gene marker, which is used as reference for OFF subpopulations. Vertical lines leave 99% of the data acquired for the non-GFP strain to the left and is used to differentiate between OFF and ON bacterial cells. Microscopy and cytometry panels show typical results of at least three independent replicates. (B) Mean GFP fluorescence intensity displayed by populations of the *fliC*::*GFP3* strain as calculated from 14 independent experiments using flow-cytometry analyses such as those displayed in A. Individual sample data is also displayed. Data sets marked with different letters display significant differences (*P* < 0.0001) as established by Tukey’s multiple comparisons test. (C) Robust coefficient of variation (RCV) obtained from flow cytometry data shown in A. Data sets marked with different letters were established as displaying significant differences (*P* < 0.05) as established by Tukey’s multiple comparisons test. (D) Confocal microscopy images of microcolonies formed within the bean leaf apoplast by a strain carrying both the *fliC*::*GFP3* gene fusion and a constitutive *eCFP* marker gene, 3 dpi by infiltration with a 10^6^ CFU/ml bacterial suspension. GFP channel shows heterogeneous expression of *fliC.* CFP channel shows constitutive expression of CFP uniformly expressed and merged panels show the combined GFP and FM4-64 channels. Red in all panels corresponds to chloroplasts autofluorescence. Scale bars correspond to 5 µm. Contrast and brightness were adjusted to improve visualization but were kept constant across different channels. (E and G) Histograms show GFP fluorescence *versus* cell count for the *fliC*::*GFP3* strain grown overnight in LB (E) or 24 hours in HIM (G) before sorting (left panels) and after sorting (right panels). Red lines in the left panels indicate the gates drawn to separate *fliC*::*GFP3* bacteria according to the level of GFP fluorescence (low or high). (F and H) Graph represents relative ratio between the halo diameters in swimming plates seeded with cells sorted according to their high expression of GFP and those sorted from the same cultures displaying low expression, shown in E. Sorting was carried out using either overnight LB-grown (F) or 24 hours HIM-grown (H) cultures. From each culture undergoing sorting, 2 µl of each of the sorted samples (low or high) adjusted to 10,000 sorted events, were inoculated into soft-agar swimming motility plates. Images were then taken 1 and 3 days after inoculation (for LB-grown sorted samples) or 3 days (for HIM-grown sorted samples). Halos obtained were measured and the ratio between the diameters of high versus low expression within each sorted pair was calculated. Data was collected from 3 (HIM-grown) to 4 (LB-grown) independent experiments each including 3 biological replicates. Asterisks indicate that the differences were established as significant by an unpaired t test (P < 0.05) and P values are indicated.

In keeping with the notion of flagella having a role in facilitating entry into the leaf and with previous transcriptomic data^3–6,8^, leaf-surface-located bacteria are mostly Flagella^ON^. Nonetheless, flagellar expression does also display fluorescence heterogeneity in this niche (**Fig. S1**). Four days post-pressure inoculation into the leaf apoplast, after bacterial growth has taken place, bacteria extracted from the apoplast display population-average expression levels significantly lower than those observed in media-grown populations (**Fig. 1B**), in agreement with previous transcriptomic data^11,12^. However, most of these bacteria are Flagella^ON^ (**Fig. 1A, lower panels**). Cell-to-cell differences in fluorescence amongst apoplast-extracted bacteria were more pronounced than in laboratory media, with a significantly larger robust coefficient of variation (RCV, **Fig. 1C**). This variability was also apparent within the apoplastic microcolonies within the plant leaf, differing from the homogeneous distribution of fluorescence displayed by constitutively expressed eCFP (**Fig. 1D**).

When bacterial populations exponentially growing in rich medium (LB) (**Fig. 1E**) were sorted by fluorescence-activated cell sorting (FACS) based on the individual cell level of GFP fluorescence, the resulting populations, enriched in bacteria expressing either low or high levels of the *fliC::GFP3* reporter (**Fig. 1E; right panels**), displayed differences in swimming motility. Although sorting can cause physical damage to the flagellar filament, potentially reducing differences in motility between the sorted subpopulations and introducing variation between independently sorted replicas, the high-to-low ratio between the halo diameters for each pair of sorted subpopulations was significantly higher than 1.0 (**Fig. 1F**). Similar results were obtained in HIM (**Fig. 1G-I**), albeit motility ratios and variation were lower than those obtained from LB cultured bacteria.

Thus, these results indicate that the flagellar system is phenotypically heterogeneous in *P. syringae* in all conditions tested. The results also show that, despite flagellin being an immune elicitor, flagella are expressed by part of the population within the apoplast.

### Flagellar and T3SS expression display counter-regulation at the single-cell level

To evaluate the relation between the expression of flagellar and T3SS genes at the single-cell level, we used a dual-reporter strain carrying transcriptional fusions of *GFP3* to T3SS *gene hrpL* and tdTomato (tdT) to the *fliC* gene. Population expression profiles for *fliC::tdT* are consistent with those observed for *fliC::GFP3*, albeit fluorescence intensity is lower for the tdT reporter than for GFP (**Fig. S2**). The dual reporter strain displayed heterogeneous expression of both systems during growth on HIM (**Fig. 2A**). Flow cytometry analysis showed the formation of two large subpopulations, formed by bacteria favoring expression of either the T3SS genes or *fliC* (T3SS^ON^/Flagella^OFF^ and T3SS^OFF^/Flagella^ON^) (**Fig. 2B; Fig. S3A**). Although these two phenotypic combinations were more abundant, bacteria displaying expression of both systems (T3SS^ON^/Flagella^ON^), or to a lesser extent none (T3SS^OFF^/Flagella^OFF^) were also detected. Apoplast-extracted bacteria (4 days after inoculation by infiltration) also displaying all phenotypic combinations, although green bacteria (T3SS^ON^/Flagella^OFF^) were more abundant than in HIM (**Fig. 2A-B; right panels**). The apoplastic population did not display the bimodal distribution of GFP seen in HIM (**Fig. 2B; Fig. S3**), and previously reported for T3SS single-cell cell expression^27^.

**Figure 2.**
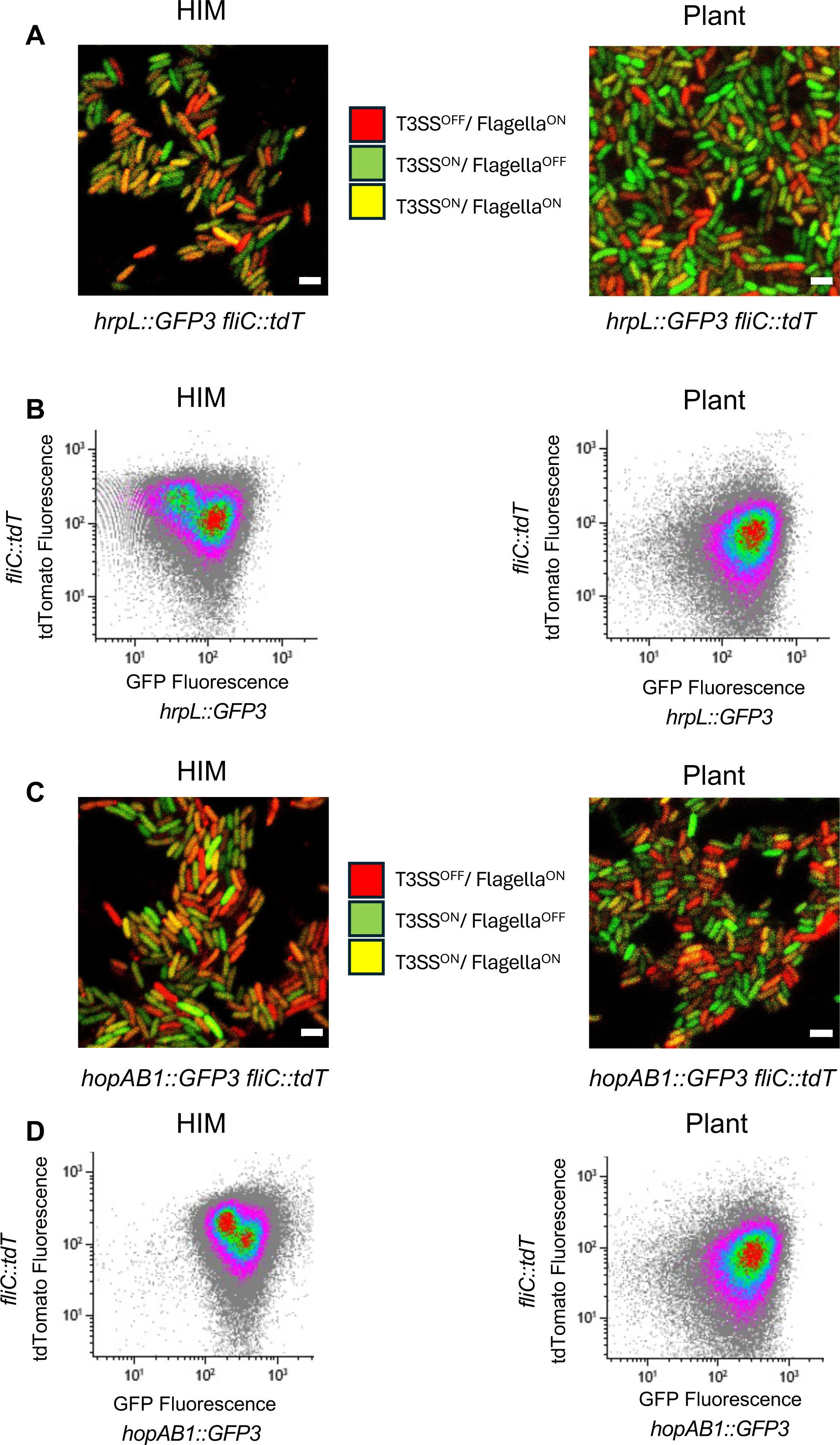
Flagellar and T3SS expression display independent heterogeneous expression at the single-cell level. (A) Confocal microscopic images of dual-reporter *hrpL*::*GFP3 fliC*::*tdT* strain grown in HIM for 24 hours (left panel) or extracted from bean leaf apoplasts 4 dpi by infiltration with 5×10^4^ CFU/ml (right panel). Scale bars in correspond to 2 μm. Contrast and brightness were adjusted to improve visualization and use throughout. (B) Flow cytometry analysis of bacteria described in A. Graphs show the fluorescence intensity of GFP *versus* that of tdTomato. Data is represented as arbitrary units and collected for 100,000 events per sample. Results shown in the figure are representative of three or more independent experiments. (C) Confocal microscopic images of dual-reporter *hopAB1*::*GFP3 fliC*::*tdT* strain grown in HIM for 24 hours (left panel) or extracted from bean leaf apoplasts 4 dpi by infiltration with 5×10^4^ CFU/ml (right panel). Scale bars in correspond to 2 μm. Contrast and brightness were adjusted to improve visualization and use throughout. (D) Flow cytometry analysis of bacteria described in C. Graphs show the fluorescence intensity of GFP *versus* that of tdTomato. Data is represented as arbitrary units and collected for 100,000 events per sample. Results shown in the figure are representative of three or more independent experiments.

Analysis of another T3SS gene, *hopAB1* (encoding a T3SS effector), which displays patterns of phenotypic heterogeneity similar to *hrpL* but with higher expression levels^27^ gave similar results both in HIM and *in planta* (**Fig. 2C-D**).

### Expression of T3SS or flagella carries growth penalties

In *Salmonella*, expression of the T3SS encoded by *Salmonella* pathogenicity island 1 (SPI1) causes severe growth delays, and this growth penalty is believed to be linked to the maintenance of phenotypic heterogeneity^18,29,30^. To test whether expression of the T3SS in *P. syringae* has an impact on bacterial fitness, we used competitive index assays (**Fig. 3A**) to compare growth between a *ΔhrpL* mutant strain and the wild type in both LB medium and HIM. CI analysis shows that the *ΔhrpL* mutant strain outgrows the wild type in HIM (CI=1.75+/-0.15) but not in rich medium (CI=1.07+/-0.08), where the T3SS genes are not expressed (**Fig. 3B**). An increased growth rate of *ΔhrpL versus* wild type was also observed in separate cultures growing in HIM (**Fig. 3C**). *In planta*, the wild type strain outgrows a derivative constitutively expressing HrpL from a plasmid (pHrpL) (CI=0.67+/-0.06), which results in overexpression of T3SS components and effectors^27^(**Fig. 3D**), further supporting the notion that expression of the T3SS carries a growth penalty.

**Figure 3.**
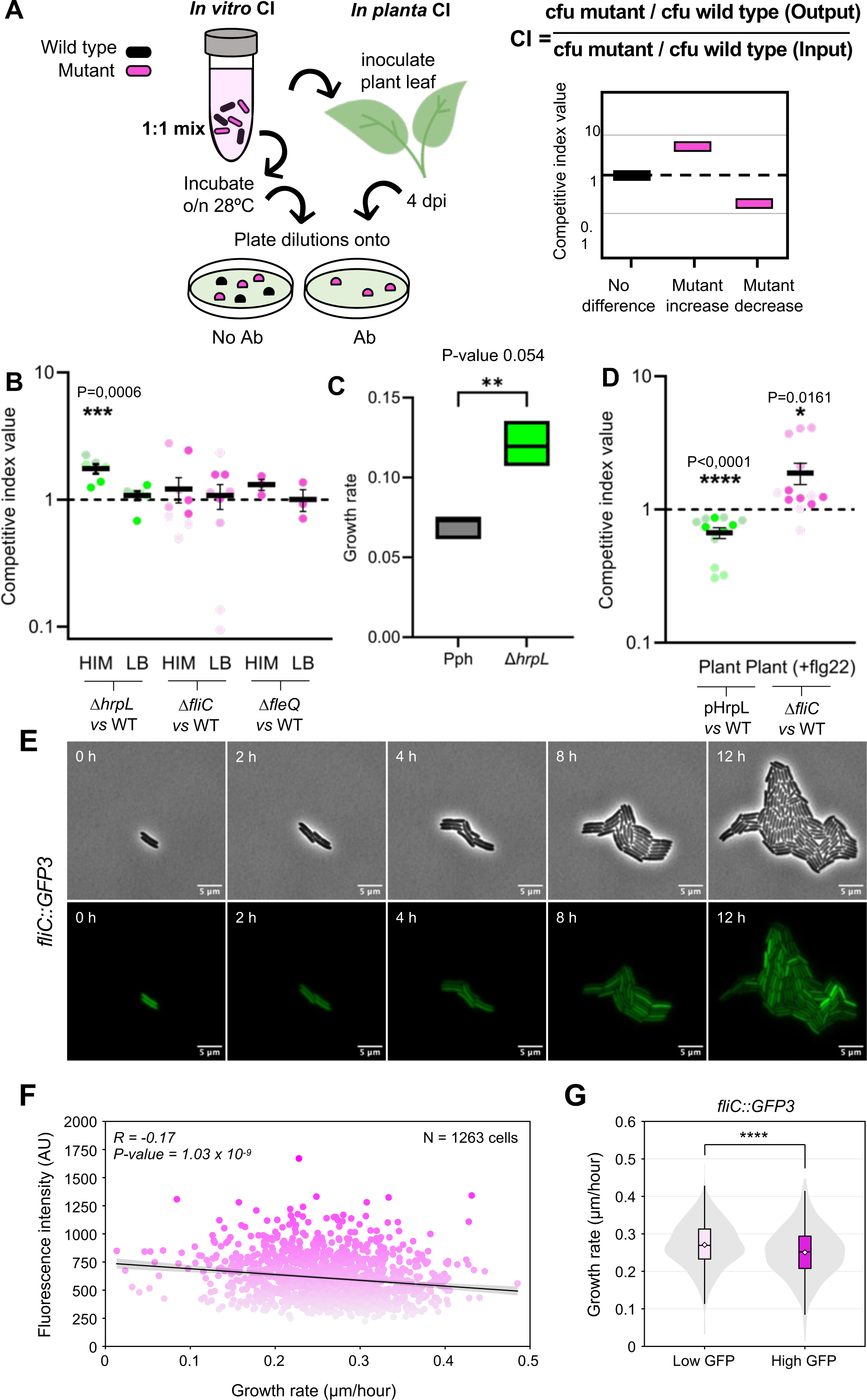
Flagellar and T3SS expression impact on bacterial growth. (A) Summary of experimental approach for competitive index (CI) analysis. First a 1:1 mixture is prepared using O/N grown LB cultures of wild type and mutant (or plasmid-carrying) strain. Then this mixture is either used to inoculate plant leaves (*in planta* CI) or to start a mixed culture in fresh medium (*in vitro* CI) which is then grown O/N at 28°C. Inoculated plants are maintained for 4 dpi before bacteria are pressure-extracted from the apoplast. O/N grown 1:1 media culture and bacterial apoplast extractions are then serially diluted and plated onto LB plates with and without antibiotic selection (only mutant bacteria grow) and wild type and mutant CFU in the output samples determined. The CI is calculated as the ratio between mutant to wild type CFU in the output sample divided by mutant to wild type CFU in the input (which should be close to 1). Once the CI is calculated is established to be statistically different from 1.0, the two strains are found to grow significantly different in the corresponding conditions (*in vitro* or *in planta*), with the mutant outgrowing the wild type strain if the CI is higher than 1.0 or *vice versa* when is lower. (B) Competitive index values for the Δ*hrpL*, Δ*fliC or* Δ*fleQ* mutant strains *versus* wild type strain (WT) after 24 hours of growth in HIM or overnight growth in LB. (C) Bacterial growth rates of Δ*hrpL* mutant and wild type strains in HIM. (D) *In planta* competitive index values for a strain constitutively expressing *hrpL* from a plasmid (pHrpL) *versus* wild type (WT), or a Δ*fliC* mutant strain *versus* wild type strain, 4 days post-inoculation. Plants used for the CI of the Δ*fliC* mutant strain *versus* wild type were pre-treated with exogenous epitope flg22 to elicit plant defence responses. (B-D) Asterisks indicate results significantly different from 1.0 as established by a non-parametric t-student test. P-values for those statistically different are indicated. Dots displaying same colour correspond to different biological replicates within one independent experiment. (E) Selected timelapse images of the *fliC*::*GFP3* strain during the microcolony development on agar pads in phase contrast (top) and GFP fluorescence (bottom) channels. Contrast and brightness were adjusted to improve visualization but were kept constant across frames of each timelapse. (F) Correlation between growth rates and fluorescence intensity of individual *fliC*::*GFP3* cells indicates that higher *fliC* expression is associated with slower growth. The shaded area shows the 95% confidence interval. (G) Comparison of cells with high and low GFP signal (determined by splitting the population into two groups using the median fluorescence intensity value) shows a significant growth decrease in cells with high flagellum expression of 6.5 % (Mann-Whitney U test, *P* = 3.6 x 10^-8^).

Flagellar expression, assembly and function has been reported to have a high energy cost in *E. coli* and *P. putida*^31–33^. However, we did not observe any statistically significant difference in growth for mutants *ΔfliC* or *ΔfleQ* (lacking the flagellar transcriptional activator FleQ) either by competitive index or in growth rates in either LB medium (CI=1.08+/-0.22 and 1.00+/-0.13, respectively) or HIM (CI=1.21+/-0.22 and 1.31+/-0.13, respectively)^34^. Interestingly, when we pretreated plants with exogenous flagellin epitope (flg22) to void differences associated to flagellin-mediated elicitation of immunity, the CI assay revealed a growth advantage for the *ΔfliC* mutant (CI=1.88+/-0.34) (**Fig. 3D**). This observation suggests that growth benefits due to the absence of flagellar expression may be specific to the environment. Constitutive expression of FleQ from pFleQ lead to a severe growth delay (**Fig. S4A**), although a pleiotropic effect could not be ruled out given the severity of this phenotype in the context of the results obtained for the flagellar mutants.

Since T3SS and flagellar expression are both highly heterogeneous, the use of population-level assays to estimate the impact of expression on bacterial fitness could be noisy and underestimate any associated growth penalties. Thus, we sought to analyze these impacts at the single-cell cell level using HIM-agar pads and time-lapse microscopy. In this experimental setting, heterogeneous gene activation of *hopAB1::GFP3* bacteria takes place too late in the experiment to allow observation of enough rounds of cell division for T3SS^ON^ bacteria for a confident quantification (**Supplemental movies S1-4**). However, heterogeneous gene activation of *fliC::GFP3* takes place earlier (**Supplemental movies S5-6**) and can thus be quantified (**Fig. 3E-G**). Single-cell time-lapse analyses show that higher *fliC* expression correlates with slower bacterial growth (**Fig. 3F**). After splitting the *fliC::GFP3* population into two using the median fluorescence intensity value, we detected a 6.5% growth penalty for the higher-than-median-fluorescence half compared to the lower (**Fig. 3G**). Controls carried out using a strain constitutively expressing eGFP shows that the growth rate for eGFP bacteria does not correlate with GFP intensity (**Fig. S4B-F**), supporting that the fitness cost detected in the high-expressing subpopulation of the *fliC::GFP3* strain is linked to high levels of *fliC* expression.

### Spatial phenotypic specialization during development of apoplastic microcolonies

The apoplastic Flagella^ON^/T3SS^OFF^ subpopulation is potentially more vulnerable to plant defenses since it elicits but cannot suppress immunity. We used propidium iodide to detect membrane-compromised (not viable) apoplast-extracted bacteria to determine if any of the phenotypic variants was subject to higher killing rates^35,36^. No significant differences were observed between the dead/live ratios of ON *versus* OFF subpopulations for either *fliC*, *hrpL*, *hrcU* or *hopAB1 genes* (**Fig. S7**). These results support that Flagella^ON^/T3SS^OFF^ cells are somehow protected from plant responses, most likely by T3SS^ON^ bacteria trans-complementing their defense suppression defect. We have previously established that while an active T3SS is essential to suppress immunity and allow bacterial multiplication in the apoplast, a null T3SS mutant may proliferate up to wild type levels when in close vicinity with wild type bacteria^37^. Such trans-complementation requires both wild type and T3SS mutant bacteria growing within the same microcolony, as homogeneous independent microcolonies within the same leaf, develop according to their respective ability to suppress or not local immune defenses^37^. Therefore, any trans-complementation between T3SS^ON^ and Flagella^ON^ T3SS^OFF^ bacteria will require the generation of apoplastic microcolonies that are phenotypically heterogeneous for both T3SS and flagellar expression. As seen above, the distribution of fluorescence in apoplastic microcolonies of *fliC:GFP3* bacteria does support that, at least for this locus, phenotypic heterogeneity arises during the development of the microcolony, rendering closely located ON/OFF variants (**Fig. 1E**). Close examination of the distribution of fluorescence from *fliC::GFP3* bacteria within apoplastic microcolonies (**Supplemental movie S7**) revealed areas where fluorescence is stronger (or weaker), suggesting a common local response to stimuli (or siblings’ inheritance of a similar expression pattern). But also revealed isolated bacteria displaying levels of GFP fluorescence strikingly different from those displayed by closely located peers, in keeping with stochastic phenotypic differences, as observed in the homogeneous environments of laboratory media.

We investigated how expression of T3SS and flagella is distributed within the microcolony during colonization of the apoplast, following leaf infiltration of a strain carrying either *hopAB1::GFP3* and *fliC::tdT* or *hrpL::GFP3* and *fliC::tdT* (**Fig. 4-5**). Stochastic heterogeneity of T3SS genes was apparent throughout the time-course experiment, while the zonal distribution of flagellar and T3SS fluorescence within the microcolonies was dynamic and changed as the infection progressed. Microcolonies observed 1 day post-inoculation (dpi) displayed heterogeneous expression of T3SS genes, but no apparent flagellar expression, indicating that flagella is mostly turned off at early stages of the infection process, in keeping with published transcriptome data^10–12^. But at later time points we found that both red (*fliC::tdT*) and green (*hopAB1::GFP3* or *hrpL::GFP3*) fluorescence displayed a zonal pattern with thoroughly overlayed heterogeneity (**Fig. 4**). In keeping with the counter regulation shown for these two systems (**Fig. 2**), zones within the microcolony with overall higher expression for the T3SS *hopAB1::GFP3* gene generally displayed lower expression of *fliC::tdT*, and *vice versa*, although orange and yellow spots (indicative of T3SS^ON^/Flagella^ON^ bacteria) were also found (**Fig. 4A-B**). Most remarkably, the distribution of T3SS^ON^ *versus* Flagella^ON^ areas within the microcolonies showed a consistent spatial pattern in relation to the nearest plant cell(s) (**Fig. 5**): the side(s) of the microcolony closest to the lower epidermis (thereby closest to their inoculation point; **Fig. 5A-C**), or to a spongy mesophile host cell (**Fig. 5A and D-E**), were predominantly green, *i.e.* T3SS^ON^, whereas the distal side of the microcolony with regards to the host cell, *i.e.* the innermost or furthermost parts of the microcolonies, were predominantly red, *i.e.* Flagella^ON^ (**Fig. 5; Supplemental movies S8-9)**. These results support the notion of cooperative virulence taking place within a spatially structured microcolony during *P. syringae* colonization of the plant apoplast, with TTSS^ON^ bacteria trans-complementing the defense suppression defect of TTSS^OFF^ bacteria (including Flagella^ON^ bacteria).

**Figure 4.**
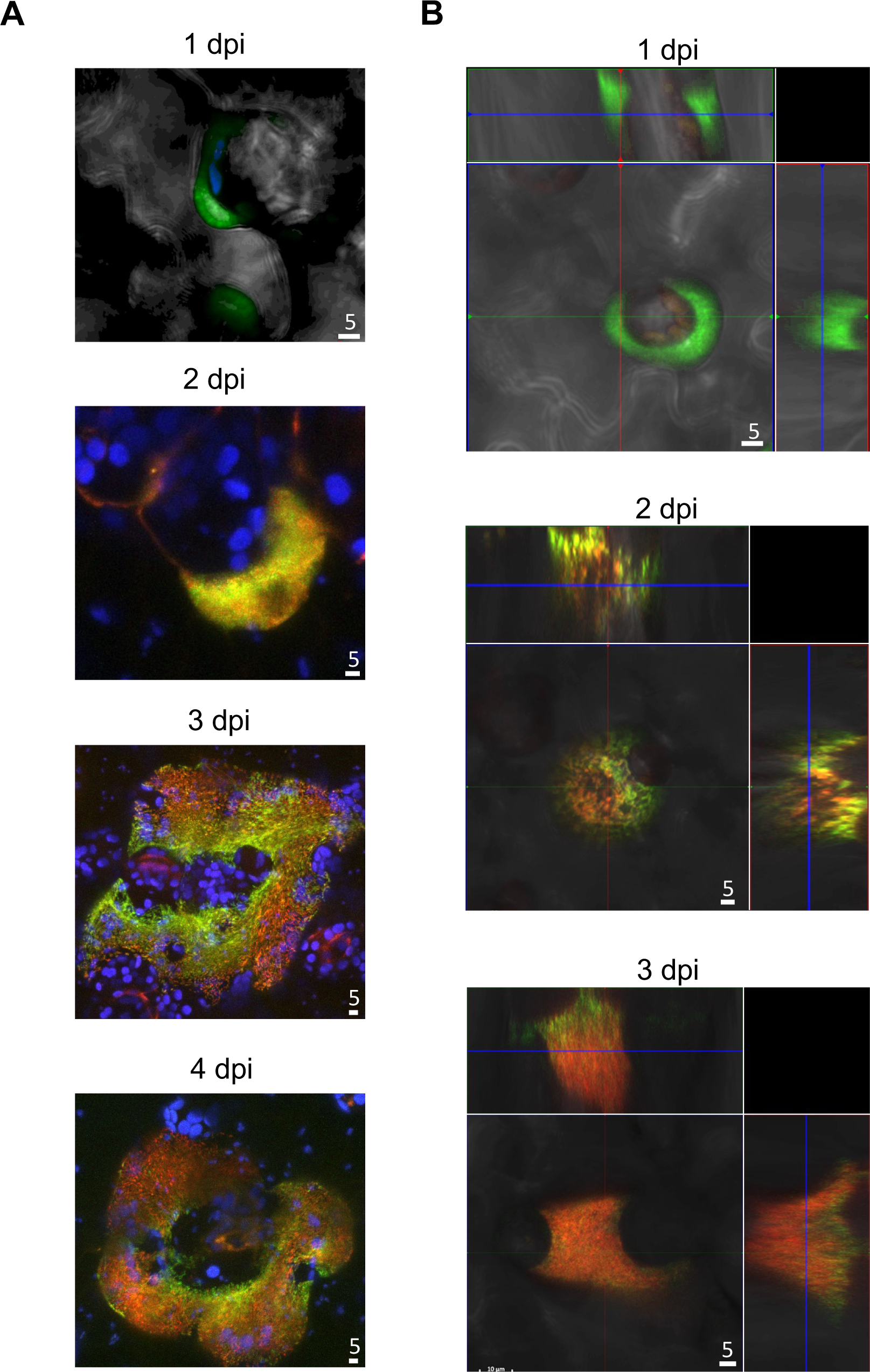
Time-course distribution of flagellar and T3SS expression during the development of apoplastic microcolonies reveals stochastic expression and different temporal dynamics for T3SS and flagella genes. Selected images of apoplastic microcolonies of the strain carrying *hopAB1*::*GFP3 fliC*::*tdT* at different days post inoculation (dpi) with 5×10^4^ CFU/ml bacterial suspensions. (A) Images correspond to Z-stack compilations at each different time points. (B) Images include orthogonal projections Scale bars correspond to 5 µm.

**Figure 5.**
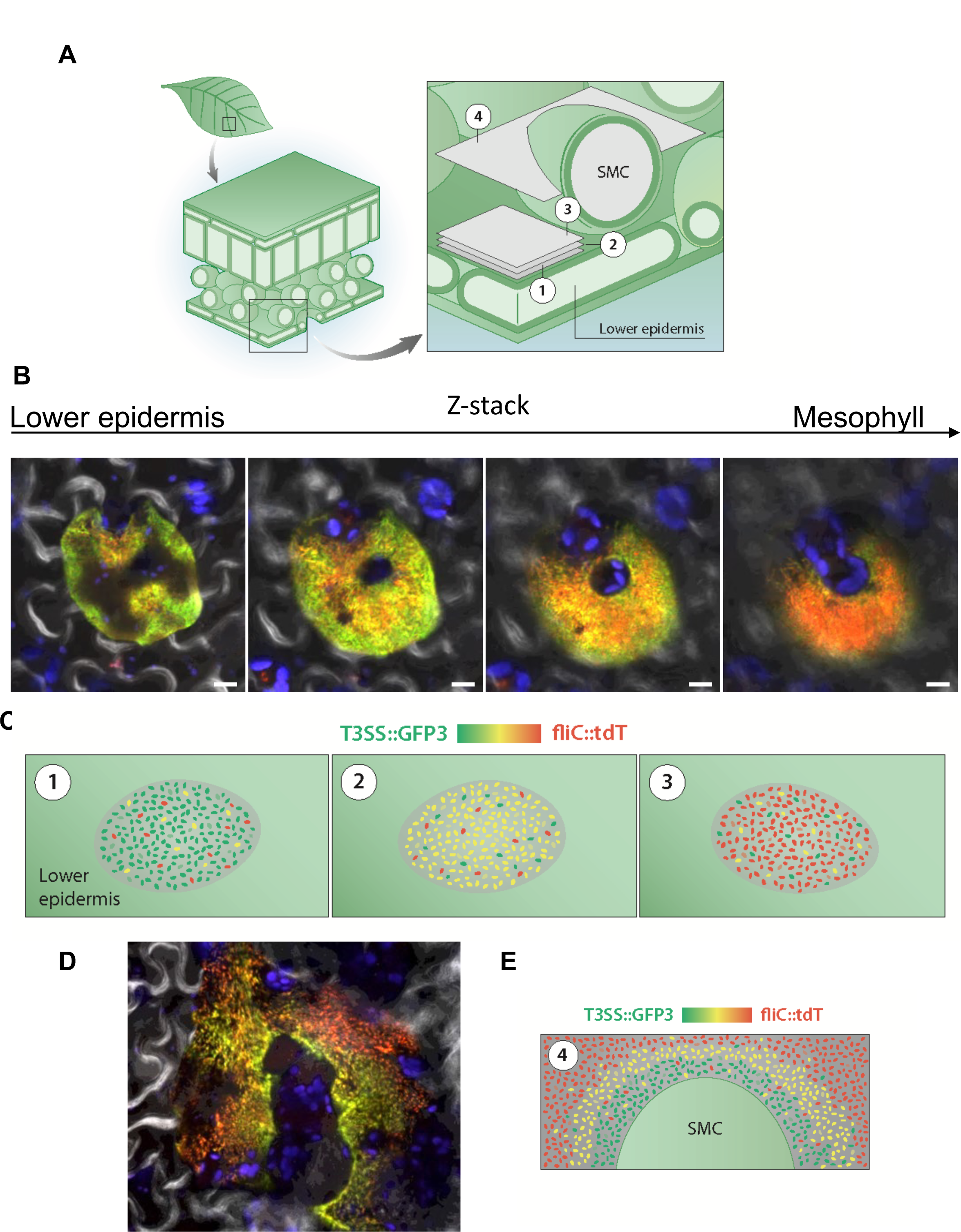
Flagellar and T3SS expression is stochastic and spatially structured within apoplastic microcolonies. (A) Schematic representation of a leaf displaying the simplified 3D leaf structure and a close up to the area where apoplastic microcolonies are detected. Position and orientation of the Z-stack acquisition shown in B and D are indicated.1-3 represent three planes of a bacterial microcolony growing from the inner surface of the lower epidermis inwards, with 1 being the closest to the cell surface and 3 the furthest (as the microcolony analyzed in the Z-stack images shown in B and supplementary movie S8. 4 represents a plane cutting through a bacterial microcolony growing wrapped around a spongy mesophyll cell (SMC) (as the microcolony and frame shown in D and the Z-stack shown in supplementary movie S9). (B) Selected images of different frames of a Z-stack acquisition taken from an apoplastic microcolony of the strain carrying *hrpL*::*GFP3 fliC*::*tdT* strain at 4 dpi with 5 x 10^5^ CFU/ml bacterial suspensions (complete Z-stack shown in supplementary movies S8). Contrast and brightness were adjusted to improve visualization but were kept constant across the different times and frames of the Z-stack acquisition. Scale bars correspond to 10 µm. Lower epidermis and mesophyll indicate the relative position of the frames within the leaf. (C) Schematic representation of the distribution of fluorescence for *fliC::tdT* and *T3SS::GFP3* within the type of apoplastic microcolonies located in A (1-3) and shown in B: Z-stack planes closest to the lower epidermis (abaxial side) showing predominantly green bacteria and those furthest predominantly red, with the intermediate planes displaying a yellow majority. Heterogeneity is also depicted within each plane. The legend indicates green as corresponding to T3SS^ON^/ Flagella^OFF^ bacteria, red to T3SS^OFF^/ Flagella^ON^ and yellow and oranges to the expression of both loci to different intensities. (D) Selected frame of a Z-stack acquisition taken from an apoplastic microcolony of the strain carrying or *hopAB1*::*GFP3 fliC*::*tdT* strain (complete Z-stack shown in supplementary movies S9). Scale bars correspond to 20 µm. (E) Schematic representation of the distribution of fluorescence for *fliC::tdT* and *T3SS::GFP3* within the type of apoplastic microcolonies located in A (4) and shown in D: bacteria closest to the cell are predominantly green, turning to yellow and red as the distance from the cell surface grows. Heterogeneity is also displayed throughout the microcolony. The legend indicates green as corresponding to T3SS^ON^/ Flagella^OFF^ bacteria, red to T3SS^OFF^/ Flagella^ON^ and yellow and oranges to the expression of both loci to different intensities.

### Active exit from infected tissues is carried out by Flagella^ON^ bacteria

The increasing numbers of Flagella^ON^ bacteria as the apoplastic microcolonies develop suggested a potential function for this phenotypic variant at later stages of the infection process. Since previous data supports flagellar motility is not required for systemic spread, we considered whether flagellar activation could be linked to the following step on the life cycle of the pathogen: leaving the infected tissue to move to a new niche or host. To address this possibility, we took leaves at 1 dpi (asymptomatic leaves) and 7 dpi (fully symptomatic leaves) and submerged them into a sterile MgCl_2_ solution for 30 mins. Then, we removed the leaves and used confocal fluorescence microscopy to evaluate single-cell cell *hopAB1::GFP3* and *fliC::tdT* expression in the bacteria present in the MgCl_2_ solution. We found that many bacteria naturally exited the leaf tissue, even from intact 1 dpi leaves showing no signs of tissue damage. We then compared the expression of T3SS and/or flagellin within the populations of bacteria either exiting the leaf by their own means or forcefully extracted from the apoplast by mechanical means (**Fig. 6**). No significant differences were found in the distribution of fluorescence for *hopAB1::GFP3* expression between naturally exiting *versus* apoplast-extracted bacteria at 1 dpi (**Fig. 6A-B**). In keeping with results shown in **Fig. 5**, apoplast-extracted bacteria at 1 dpi showed a vast majority of Flagella^OFF^/T3SS^ON^ and almost no Flagella^ON^/T3SS^OFF^ bacteria (**Fig. 6A upper panel**). In contrast, bacteria naturally leaving leaves at 1 dpi were significantly enriched in Flagella^ON^ *versus* those present in apoplast-extracted (**Fig. 6A and B**), indicating that Flagella^ON^ bacteria are more efficient in actively exiting the leaf in natural conditions. These results support that bacterial exit from asymptomatic tissues is flagella-dependent.

**Figure 6.**
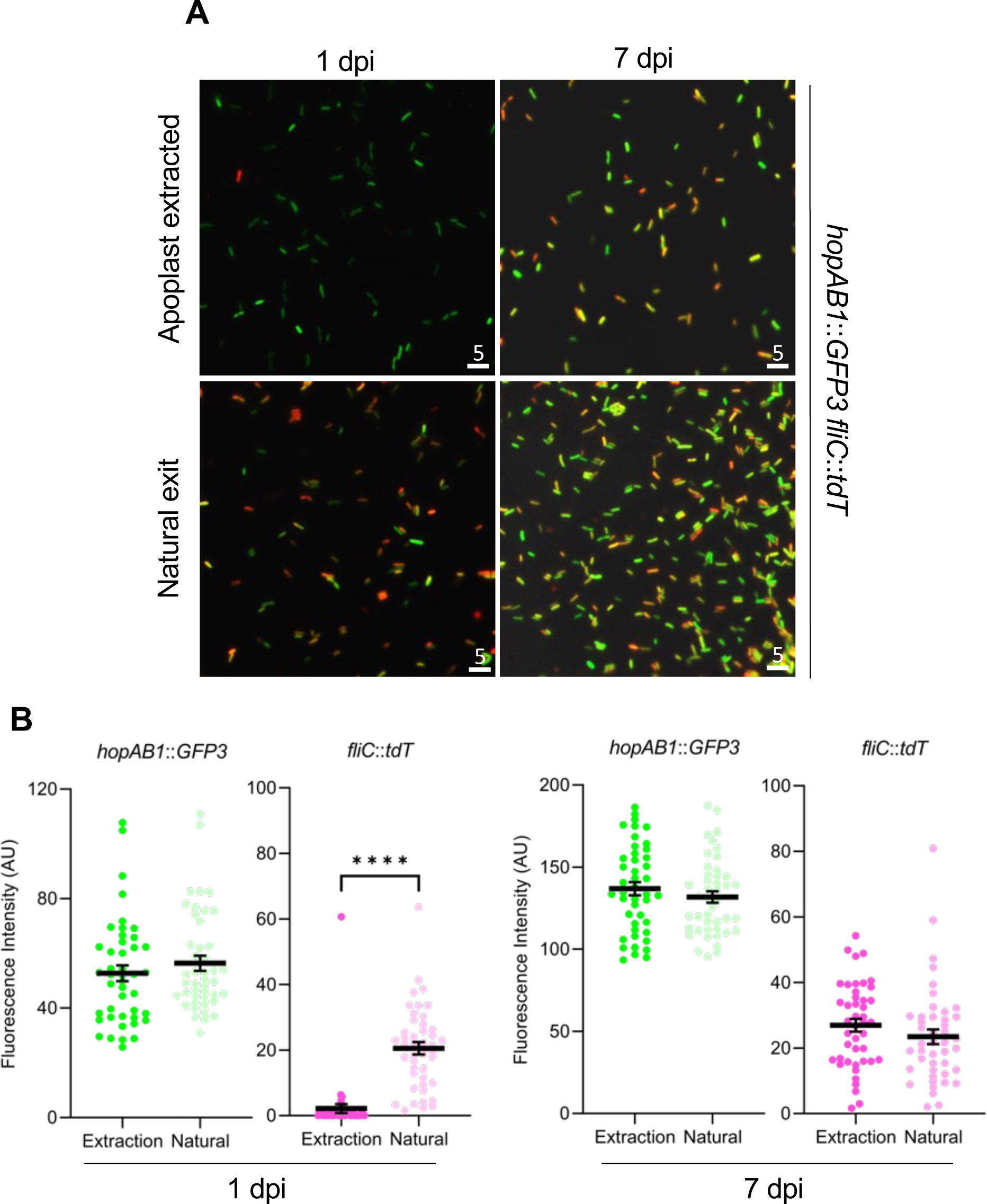
Active exit from infected tissues is carried out by Flagella^ON^ bacteria. (A) Selected images of apoplast-extracted bacteria (using negative pressure; upper panels) or bacteria exiting the tissue on their own (during leaf incubation within a MgCl_2_ solution; natural exit, lower panels) at 1 dpi (asymptomatic tissue) or 7 dpi (fully symptomatic necrotic tissue). Contrast and brightness were adjusted to improve visualization but were kept constant across the different conditions. (B and C) Fluorescence quantification of the images obtained in (A) using Fiji software. Graph shows arbitrary units of GFP fluorescence, corresponding to the expression of the T3SS gene fusion *hopAB1::GFP3* in the samples, or tdTomato fluorescence corresponding to the expression of *fliC*::*tdT*. Each dot corresponds to an individual bacterium as analyzed from the image. Comparisons between apoplast extraction and natural exit were carried out per each sample using an unpaired t test (P<0.0001) and the results shown for each pair.

This bias towards Flagella^ON^ among naturally exiting bacteria was no longer observed when the analysis was carried out using 7 dpi fully symptomatic leaves, where leaf integrity has already been lost due to the onset of necrosis. Based on these results, we propose that flagellar motility is not only important for bacterial entry into the leaf but also for rapid and active exit of apoplast-growing populations prior to tissue collapse.

## DISCUSSION

Bacterial encounters with changes in the environment lead to the activation of specific genes, which is often assumed to take place in a nonfluctuating and homogeneous manner. However, such studies are often carried out by establishing average values for entire populations, losing out potential cell-to-cell variation^38^. In the last decades, an increasing number of bacterial traits have been shown to respond differently across clonal populations to environmental changes and/or complex niches. These response variations may reflect differences in their microenvironment but can also be caused by stochastic cellular changes (gene expression noise) in particular regulatory circuits (which enhance of even exploit noise) leading to phenotypic heterogeneity.

### Flagellar phenotypic heterogeneity

*P. syringae* produce 1-5 polar flagella^39,40^ upon activation of FleQ, the transcriptional flagellar regulator in *Pseudomonas*^21,22,41^, under a regulatory cascade mostly uncharacterized. Conservation of gene arrangement and promoter motifs with *P. putida,* which produces 5-7 polar flagella^21,42^, supports a similar organization for *P. syringae*. Nonetheless, our results show that the *P. syringae* flagellar regulation also shares an important feature with that of peritrichous *Salmonella enterica*: stochastic cell-to-cell variation (**Fig. 1; Fig. S1-4**)^13–19,43^. Whether such phenotypic variation extends to other pathogenic or commensal Pseudomonas remains to be investigated. Bistable activation of the *Salmonella* flagellar system is established from a noisy heterogeneous expression pattern through two independent mechanisms: (*i*) a double-negative feedback loop involving RflP and FliZ regulators, governing bistability of class 2 genes, and (*ii*) a developmental checkpoint established by σ28-FlgM driving bimodal expression of class 3 genes^13–16,20,43^. In *P. syringae*, a dual mechanism is also involved in turning T3SS noisy expression into bimodality^27^: (*i*) a double-negative feedback loop (involving HrpV, a repressor of HrpS that reduces HrpL levels, and HrpG an anti-repressor^25,44^, and (*ii*) a positive feedback loop by the main component of the Hrp T3SS secretion pilus HrpA^39,45^. Future research into the regulatory elements governing flagellar synthesis in *P. syringae* would help to identify candidates potentially involved in establishing phenotypic heterogeneity in flagella.

Flagellar heterogeneity in *P. syringae* was observed in all conditions tested, including leaf surface and apoplast (**Fig. 1 and S2**), with occasional bistability in media grown populations. Average mean expression in these two locations fit those reported in transcriptomic studies^10–12^. However, it was unexpected to find such a large percentage of surface-localized bacteria not expressing flagella, and particularly so to find so many apoplast-localized bacteria expressing high levels of flagella at late stages of the infection (**Fig. 1 and 5**).

### Crosstalk and heterogeneous expression as adaptations to alleviate fitness costs

Negative crosstalk between T3SS systems and flagella (**Fig. 2-3**) has been shown for other bacterial pathogens. In *Salmonella*, flagellar expression is counter regulated with the T3SS encoded by SPI1 (*Salmonella* encodes two functionally separate T3SS^46–48;^ which as flagella in this species displays phenotypic heterogeneity and bistable expression and is associated to a severe growth penalty^13–16,30,49,50^. In *P. aeruginosa*, the global regulator GacA has been proposed as linked to flagella-T3SS counter regulation ^51^, whereas in plant pathogenic *Erwinia amylovora*^52^, it has been reported to be carried out by HrpL. Twenty-two transcription factors (TF) regulate expression of the T3SS in *P. syringae*, seven of which, including GacA^26,53–56^, have been also reported to affect flagellar expression^57^, thus providing several candidates to mediate T3SS-flagella counter regulation in this pathogen. Flagellar genes were reported to be negatively regulated by T3SS regulators HrpRS (two T3SS-specific enhancer-binding proteins that act as a hetero hexamer activating transcription from the *hrpL* promoter^58^; but not by HrpL^59^. However, since overexpression of HrpL impacts average flagellar expression (**Fig. 3**), either HrpL directly represses flagellar expression or elements downstream of HrpL feed back into the regulatory cascade(s) established by the upstream regulator(s) repressing flagellar expression.

In any case, variation in the expression of the flagellar systems was not significantly impacted by overexpression of HrpL (**Fig. 3**), supporting that phenotypic heterogeneity in this system arises through an independent mechanism from the mechanism driving bistablilty of the T3SS^27^. Independent switching of SPI1 T3SS and flagella also takes place in *Salmonella*^60^.

Despite examples of counter-regulation between T3SS and flagella that accompany our own, these two systems can be coordinately expressed in other cases, *i*.*e*. in enteropathogenic *E. coli* (EPEC) flagellar assembly requires a functional T3SS^61^. Different scenarios might be associated to differences in the metabolic costs and growth penalties of the systems involved. The metabolic cost of flagellar motility has been demonstrated for *E. coli* and *P. putida*^31–33^. We have found that expression of the flagellar system also has a fitness cost in *P. syringae* (**Fig. 3 and S5**). Flagellar motility costs are clearer at the single-cell cell level (**Fig. 5**) and may vary depending on the conditions, as results *in planta* showed a significant cost at the population level not detected in media. Flagellar motility in *P. syringae* may have higher fitness costs in other environments or under stress conditions, as reported for *P. putida*^32^. In *Salmonella*, the strong metabolic costs linked to expression of the SPI1 T3SS has been proposed to be a factor favoring both its counter regulation with flagella and the maintenance of its phenotypic heterogeneity^14,29,30,49,50^. Although not as severe as that reported for SPI1 in *Salmonella*, we have found evidence of a growth penalty associated to T3SS expression in *P. syringae*: (*i*) during growth in HIM and (*ii*) *in planta* (**Fig. 3**), although in the latter case constitutive expression of HrpL hindering heterogeneity could be a confounding factor.

Phenotypic heterogeneity is considered particularly advantageous when affecting loci involved in producing immunogenic and/or energetically costly goods, with the fitness cost to the individual producer cells and the corresponding benefit to the population as a whole defining a cooperative behavior^29,30,60,62–70^. The growth penalties detected for flagellar and T3SS expression in *P. syringae* could provide a selective advantage to phenotypically heterogeneous populations and thus determine selection of such a trait (phenotypic heterogeneity of each of these systems) and of the underlying genetic and/or epigenetic mechanism(s). In addition, for bacteria engaging in host interactions, a heterogeneous pattern of expression that lowers the overall amount of flagellin displayed by the bacterial population, as a whole or by a microcolony at a local level, could provide additional advantages by reducing defense elicitation and/ or facilitating immune suppression.

These results extend the link between virulence factor production and slow growth reported for some virulence traits in human pathogens to plant pathogens^70^, and as such support the notion of cooperative virulence in *P. syringae*. Additionally, cooperative virulence has been proposed in animal pathosystems to provide additional advantages by preventing the rise of nonproducing mutant variants at the expense of the producers (so called cheaters)^66,70^. Phenotypically OFF bacteria can outcompete less frequent mutant OFF variants and safely revert to ON, thus preventing loss of the function for the population and stabilizing the cooperative behavior^66^. Cheater mutant variants affected in type III secretion have been reported to rise in *P. syringae* populations during colonization of *Arabidopsis thaliana*, however at frequencies lower than expected considering the potential for exploitation of T3SS effectors as public goods^37,71^. Whether cooperative virulence through phenotypic heterogeneity helps to limit the rise of such variants in *P. syringae* may be difficult to establish, but is a plausible hypothesis given the results presented here.

### Adaptive value of phenotypic heterogeneity of T3SS and flagella *in planta*

Phenotypic heterogeneity can provide a strategy to cope with rapidly changing and/or fluctuating environments, with preexisting subpopulations already adapted to incoming stresses which can overcome these faster^72,73^. Such ‘bet-hedging’ (or ‘risk-spreading’) strategies guarantee that the population will contain a fitter subpopulation to allow the survival of the genotype in case of rapid change(s) in the environment^74^. Phenotypic heterogeneity can also have adaptive value as a ‘division of labor’ strategy when a bacterial population that diversifies allows the distribution of tasks among phenotypically different subpopulations that thus cooperate benefitting the entire population, such as discussed above in the case of metabolically costly traits^65^. While in bet-hedging, one subpopulation is fitter than the other in each environment, division of labor increases the fitness of the entire population^75^. These two strategies are not mechanistically exclusive and can overlap^76^. We have found no evidence supporting that any of the subpopulations generated through T3SS and flagellar expression heterogeneity is specifically benefiting during plant colonization, as would be expected from a proper bet-hedging scenario. However, different defense elicitation contexts, such as colonization of resistant plant hosts, might differentially benefit certain phenotypic variants, as shown for variants generated through genomic reorganizations in *P. syringae*^77–80^. In addition, an analysis of the broader life cycle of the pathogen, where the fate of the pathogen beyond the initial infection is included (see discussion below) could potentially support a bet-hedging scenario. In any case, results so far support a division of labor scenario, with T3SS^ON^ bacteria complementing T3SS^OFF^ likely through suppression of immunity by ‘common goods’ T3SS effectors (**Fig. 6, Fig. S5,** ^37^). The percentage of T3SS^ON^ bacteria required for effective *trans* complementation of T3SS^OFF^^37^ within a heterogeneous population, particularly within a context where part of the population is expressing immunogenic flagellin, is probably important and is likely to be counter balanced by growth trade-offs.

### Phenotypic heterogeneity and spatial differentiation within apoplastic microcolonies

As different eukaryotic cell types work together within tissues of higher organisms, cooperation between bacterial populations emerges within spatially organized structures^76^. However, such spatially structured cooperation has been rarely shown within the context of disease^81^. Indeed, although phenotypic heterogeneity is well documented in planktonic cultures, the extent to which similar responses occur in spatially structured communities is less understood^82^.

In apoplast-extracted bacteria, expression of T3SS genes display heterogeneity but not bistability as observed during growth within inducing medium^27^. The regulatory loops involved in establishing bistable expression of the T3SS in HIM^27^ are in place during colonization of the apoplast^26,83^. However, the apoplast is not homogeneous and apoplastic bacteria are expected to encounter different stimuli specific to their microenvironment, which in addition can change over time. These potentially lead to cell-to-cell variation orthogonal to that generated through phenotypic heterogeneity for the two loci under study. Indeed, time course expression analysis of T3SS and flagellar expression during plant colonization shows phenotypic heterogeneity at all time points and bacterial locations, but also an overlapping spatially structured pattern (**Fig. 7**). On the surface of the leaf, the vast majority of the population is T3SS^OFF^ regardless of the single-cell cell status of flagellar expression, in keeping with transcriptome data ^10^. Once within the apoplast, the bacterial population changes to a majority of T3SS^ON^/ Flagella^OFF^, with clear stochastic heterogeneity for the T3SS (**Fig. 5; 1 dpi**). From 2 dpi onwards, heterogeneity is the norm for both loci, but areas predominantly T3SS^ON^ or Flagella^ON^ within the microcolonies appear. These areas show a clear three-dimensional pattern: areas closely associated to host cell surfaces are enriched in T3SS^ON^ bacteria, whereas distal areas from the host cell surface, or indeed the innermost part of the microcolony, are predominantly Flagella^ON^ (**Fig. 5 and 7; Supplemental movies S4-5**). This is consistent with the notion of T3SS^ON^ bacteria being more abundant where T3SS-mediated activity is relevant for the interaction with the host cell and explains the lack of selective killing of T3SS^OFF^ bacteria observed by dead/live staining^27^. Such phenotypic differentiation of a clonal population into spatially distributed subpopulations that cooperate in a complex more natural environment (*i*.*e*. different from laboratory media growth settings) has rarely been investigated, particularly for more than a single-cell trait at once^84^, but it is reasonable to assume that is likely to take place in many natural settings, particularly in the context of host colonization processes. Spatially structured cooperation in the context of disease have been demonstrated for the cholera toxin and toxin-coregulated pilus in *Vibrio cholerae*, and for the nitric oxide (NO)-detoxifying *hmp* gene in *Yersinia pseudotuberculosis*, in intestinal and spleen mice microcolonies respectively^81,85^. Spatial heterogeneity has also been described through single-cell omics within biofilms of another human pathogen, *P. aeruginosa*, which included variation on flagellar expression^86^.

**Figure 7.**
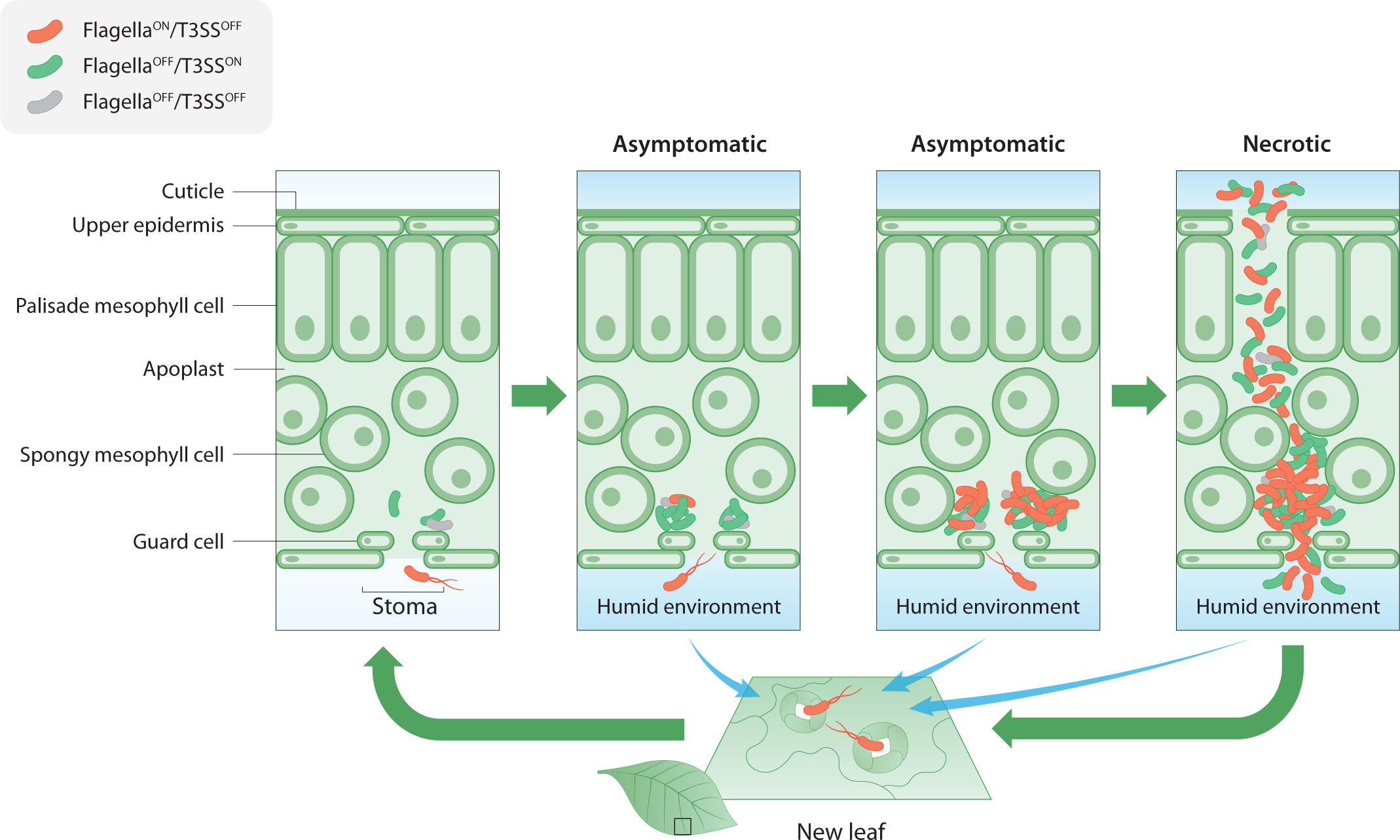
T3SS and flagella phenotypic heterogeneity overlays with dynamic and spatial expression patterns during interaction with the plant host. Model summarizes the dynamic of expression of these two loci during *in planta* growth. Starts at the beginning of the infection (top left). A leaf section with different structural elements indicated, illustrates entry of swimming bacteria into the apoplast through a stoma, where bacteria expression profiles switchs from Flagella^ON^/T3SS^OFF^ to to start the interaction with host-cell needed to suppress defenses and to allow bacterial multiplication. During the initial multiplication (1 dpi) bacteria maintain a predominant Flagella^OFF^/T3SS^ON^ expression profile with orthogonal stochastic switching to T3SS^ON^, regardless of flagellar expression which is reduced to a very limited proportion of the population. However, Flagella^ON^ bacteria are particularly capable of exiting the leaf within a humid environment and the exiting population is enriched in Flagella^ON^ bacteria. As the microcolony grows those bacteria closest to the host cell remain T3SS^ON^ with the occasional stochastic switch to T3SS^OFF^, but the growing side of the microcolony, father away from the host cell switch to Flagella^ON^/T3SS^OFF^. The latter also show overlaid stochastic heterogeneity. When the infection reaches the last stages and the plant tissue becomes necrotic, bacteria of all phenotypic combinations exit the compromised tissue in wet environments. Bacteria exiting the tissues at all the stages of the infection can potentially move onto a new niche or host. Red bacteria indicate Flagella^ON^/T3SS^OFF^ bacteria, and green bacteria Flagella^OFF^/T3SS^ON,^ although bacteria expressing both to different levels can also be found. Grey bacteria indicate stochastically OFF bacteria for either locus.

Host cell signals but also local cell-to-cell interactions or lineage history^87^ could be involved in the predominant activation of the T3SS in host-proximal areas of the microcolony. How environmental drivers combine with stochastic heterogeneity to generate phenotypic variation is a current topic of interest^84^, and usually a particularly challenging one to study beyond typical *in vitro* systems if positional information is to be obtained^82^. As the apoplastic microcolonies develop, an increasing number of bacteria favors flagellar expression. A plausible explanation for this trend would be bacteria getting ready for the next step on the infection cycle: leaving the compromised tissue to colonize a new host or niche. This notion is supported by the results showing Flagella^ON^ bacteria actively exiting colonized uncompromised tissue prior to the onset of necrosis (**Fig. 6**). Very little research has been carried out on the final stages of the infection or analyzed how bacteria leave the host after colonization, so whether this active exit is biologically more relevant than passive exit from necrotic tissue remains to be explored, but *a priori* it would appear a safer strategy since host necrosis leads to the release of potentially toxic compounds. Such a strategy, could thus be considered a sort of preadaptative heterogeneity, in which a subpopulation expresses fitness- and virulence-relevant factors, which are not useful in the current, but in the following host niche(s), a proposed version of bet-hedging for microbial pathogens in which the pathogen can use environmental cues from the current niche to anticipate and preadapt a subgroup of cells for the next stages of the infection process^88^.

Only a handful of studies have analyzed expression of two bistable or heterogeneous loci simultaneously^60,89^. Ours addresses the study of two loci of relevance for bacterial-host interaction and does it on the context of host colonization, within a spatially structured dynamic microcolony, providing a comprehensive view of how processes driving bacterial variation play out in the context of a complex and changing environment of biological and applied relevance. It also provides additional examples in which division of labor benefits the population, helping to validate the importance of phenotypic heterogeneity in nature and shifting the emphasis from a mechanistic understanding of phenotypic heterogeneity to the ecological benefits and biological relevance. This study also establishes phenotypic heterogeneity and cooperative virulence as a conserved strategy by which bacterial pathogens cope with the fluctuating challenging conditions of both plant and animal hosts. Unravelling these processes underlying bacterial specialization during host colonization may provide opportunities to predict or interfere with them to prevent or hinder the progress of bacterial diseases.

## ACKNOWLEDGMENTS

We wish to thank Adela Zumaquero and Pablo García Vallejo for technical assistance and Inmaculada Ortiz-Martín for her contribution to preliminary experiments. This work was supported by project grants from Ministerio de Ciencia, Innovación y Universidades (MCIU, Spain, RTI2018-095069-B-100 and PID2021-127245OB-I00) awarded to CRB and JRA) and “ERDP A way of making Europe”. NLP received funding as a Short-Term Training Mission from COST ACTION SUSTAIN FA1208, supported by COST (European Cooperation in Science and Technology), FEMS research and training grant, and EMBO Scientific Exchange Grant. JSR was supported by Plan Andaluz de Investigación, Desarrollo e Innovación (PAIDI 2020). JSL and SVV were supported by the Swiss National Science Foundation (SNSF) through the Swiss National Centre of Competence in Research (NCCR) Microbiomes and through an Ambizione Fellowship to SVV (grant number: PZ00P3_202186). LA is funded by Aix-Marseille University (AMU) and the Centre National de la Recherche Scientifique (CNRS). MASR was supported by PID2020-116995RB-I00 funded by MCIN/AEI/ 10.13039/5011100011033 and, as appropriate, by “ERDF A way of making Europe”.

## AUTHOR CONTRIBUTIONS

Conceptualization, N.L.P., J.R.A. and C.B.L.; methodology, software and analysis, N.L.P., J.S.R., L.A., M.S.R., J.L. and S.V.V.; resources, N.L.P., J.S.R., J.L., M.S.R.; writing-original draft, C.B.L.; writing-review and editing, C.B.L., N.L.P., J.S.R., J.L., M.S.R., L.A., S.V.V., J.R.A.; funding acquisition, C.B.L. J.R.A., M.S.R., L.A., S.V.V.; all authors discussed the results and approved the manuscript.

## DECLARATION OF INTERESTS

The authors declare no competing interests.

## INCLUSION AND DIVERSITY STATEMENT

We support inclusive, diverse, and equitable conduct of research.

## METHODS

## KEY RESOURCES TABLE

## RESOURCE AVAILABILITY

### Lead contact

Further information and requests for resources and reagents should be directed to and will be fulfilled by the lead contact, Carmen R. Beuzón.

### Materials availability

This study did not generate new unique reagent.

### Data and materials availability

Time-lapse data analysis was performed using custom Python scripts adapted from Kaczmarczyk *et al.* (2022)^99^ (available here: https://github.com/JLuneau/Pseudomonas_AgarPads_fliC).

This paper does not report original code.

Any additional information required to reanalyze the data reported in this work is available from the lead contact upon request.

## EXPERIMENTAL MODEL AND SUBJECT DETAILS

### Bacterial strains and growth conditions

Bacterial strains used and generated in this work are detailed in the key resource table. *E. coli and P. syringae* strains were grown with aeration in Lysogeny Broth (LB) medium^90^ at 37°C for *E. coli* or 28°C for *P. syringae*. Antibiotics were used, when necessary, at the following concentration: ampicillin (Amp), 100 μg/ml for *E. coli* and 500 μg/ml for *P. syringae*; kanamycin (Km), 50 μg/ml for *E. coli* and 15 μg/ml for *P. syringae* derivative strains; gentamycin (Gm), 10 μg/ml; nitrofurantoin 40 μg/ml, and cycloheximide, 2 μg/ml.

To induce the expression of the *hrp/hrc* genes, bacteria were initially cultured overnight in LB at 28°C, supplemented with the appropriate antibiotic, then, washed twice in 10 mM MgCl_2_ before being cultured in *hrp*-inducing minimal medium (HIM), containing 10 mM fructose^91^. For this study, pH of HIM was adjusted to 7.0 with 10N NaOH. The initial OD was adjusted to 0.13 and cultures were incubated at 28°C with agitation.

### Fluorescent labelling of bacterial strains

Bacterial strains carrying a chromosome-located transcriptional fusion of *fliC* gene to a promoterless *tdTomato* gene was generated using an adaptation of the method previously described in Zumaquero *et al*.^92^. For the generation of the allelic exchange plasmid, two fragments of approximately 500 pb were amplified from *Pph* 1448A genomic DNA using Q5 High-Fidelity DNA Polymerase (New England Biolabs, USA); one of these fragments (A) encompasses the 3’ end of the ORF, including the STOP codon, while the other fragment (B) covers the sequence immediately downstream to the STOP codon. All primers used are listed in the key resource table. Each reaction was carried out at 98°C for 1 minute for the initial denaturation step, followed by 30 cycles at 98°C for 30 seconds, annealing at 62°C for 30 seconds, and extension at 72°C for 30 seconds, followed by 5 minutes at 72°C for the final extension step. The reaction mixture for each PCR included 0.64 mM deoxynucleotide triphosphate (dNTP) mix, 0.4 ng of each primer, 1 ng of genomic DNA, the appropriate enzyme buffer, and commercial ultrapure water (Nalgene, Rochester, NY, USA). Two μl of each gel-purified PCR product was employed as template for the subsequent fusion PCR, employing primers A1 and B2, in a PCR reaction conducted under the conditions described, with an extended elongation time of 1 min. The resulting bands, comprising the end of each ORF and its downstream sequence separated by an *EcoRV* restriction site, were A/T cloned into pGEM-T (Promega, USA) and subjected to full sequencing to discard clones carrying mutations. This process rendered the pGT-AB-*fliC* plasmid needed for generating the allelic exchange plasmid. The sequence of the promoterless *tdTomato* (*tdT*) gene was acquired through PCR amplification from the tdTomato-pBAD plasmid (Michael Davidson & Nathan Shaner & Roger Tsien, addgene plasmid #54856^93^) using the ProtFluorF and ProtFluorR primers. This PCR generated a *tdTomato* fragment with a 5’EcoRV restriction site and a 3’EcoRI restriction site. Similarly, the kanamycin cassette was amplified using the P1 EcoRV and P2 EcoRI primers, and plasmid pKD4^94^ served as DNA template to obtain the FRT-*nptII*-FRT fragment needed for the resistance to kanamycin. This fragment featured a 5’ EcoRV and 3’EcoRI restriction site. Both amplification reactions were performed using Q5 High-Fidelity DNA Polymerase (New England Biolabs, USA) in a 50 µL PCR reaction mixture consisting of 500 ng of plasmid as DNA template, 0.5 µM of each primer, 0.2 mM dNTP’s and 0.5 µL of Q5 High-Fidelity DNA Polymerase. Each reaction followed a thermal cycling protocol that began with an initial step of 98⁰C for 2 minutes, followed by 30 cycles at 98⁰C for 10 seconds, 60⁰C for 30 seconds, and 72⁰C for 45 seconds, concluding with a final extension at 72⁰C for 5 minutes. The gel-purified fragments, both the *tdTomato* and the FRT-*nptII*-FRT, underwent digestion with EcoRI and EcoRV enzymes and were cloned into the EcoRV*-*digested pGT-AB-*fliC* plasmid, rendering the allelic exchange plasmid named pGT-*fliC*::*tdT*.

Strain 1448A *fliC*::*tdT* was obtained by introducing the pGT-*fliC*::*tdT* plasmid into *P. syringae* pv. phaseolicola 1448A through electroporation, following the method described in Zumaquero *et al*.^92^. Selection was carried out on LB plates with kanamycin. Subsequently, the resulting colonies were replicated onto LB plates with ampicillin (500 μg/ml) to discard the colonies with plasmid integration, which is indicative of a single recombination event. The colonies that exhibited kanamycin resistance but ampicillin sensitivity were then confirmed through PCR, using the A1 *fliC* and B2 *fliC* primers; and through Southern blot analysis, employing the *nptII* gene as a probe, to confirm proper allelic exchange resulting from a double recombination event occurring at a unique position within the genome. To generate strains carrying two chromosomal transcriptional fusions, the pGT-*fliC*::*tdT* plasmid was transformed into the previously generated strains 1448A *hrpL*::*GFP3*, 1448A *hopAB1*::*GFP3* and 1448A *hrcU*::*GFP3* and to generate the 1448A *hrpL::GFP3 fliC*::*tdT*, 1448A *hopAB1*::*GFP3 fliC*::*tdT* and 1448A *hrcU*::*GFP3 fliC*::*tdT* strains.

Bacterial strains carrying a chromosome-located transcriptional fusion of *fliC* gene to a promoterless *GFP3* gene, 1448A *fliC*::*GFP3* strain, were generated following the method described by Rufián *et al*., 2018a^95^ with some modifications. The *GFP3-*FRT*-nptII*-FRT fragment was obtained by digesting the plasmid pGT-GFP^+^^95^ with the *EcoRI* digestion enzyme. This fragment consists of the promoterless *GFP3* gene, complete with its ribosomal binding site (RBS), followed by the kanamycin resistance gene (*nptII*), flanked by FRT (flipase recognition targets) sites, with the entire construct bordered by two *EcoRI* restriction sites. The *GFP3-*FRT*-nptII*-FRT fragment was then blunt-ended through a PCR procedure and ligated into *EcoRV*-digested pGT-AB-*fliC* through blunt-end ligation, leading to the generation of the pGT-*fliC*::*GFP3* plasmid. Subsequently, the resulting plasmid was transformed into *P. syringae* pv. phaseolicola 1448A to generate the 1448A *fliC*::*GFP3* strain.

Additionally, the constitutively-expressed fluorescent reporter gene *eCFP* was introduced into the chromosome of the 1448A *fliC*::*GFP3* and 1448A *hopAB1*::*GFP3 fliC*::*tdT* strains using a Tn*7* delivery system, as previously described by Lambertsen *et al*.^97^ to generate the 1448A *fliC*::*GFP3 eCFP* and 1448A *hopAB1*::*GFP3 fliC*::*tdT eCFP* strains.

### Generation of mutant bacterial strains

The bacterial strain carrying a deletion of the *fleQ* gene was generated following the method described in Zumaquero *et al*.^92^, which involves the generation of gene knockouts by allelic exchange, replacing the specific ORF by a kanamycin cassette. The allelic exchange plasmid pGT-Δ*fleQ* was generated as previously described for the generation of the pGT-AB-*fliC* using primers A1 Δ*fleQ*, A2 Δ*fleQ*, B1 Δ*fleQ* and B2 Δ*fleQ* and the same experimental settings described above. The FRT*-nptII*-FRT fragment was obtained by PCR amplification using P1 EcoRI and P2 EcoRI primers and pKD4 as template, and the kanamycin cassette was finally inserted by ligation in EcoRI restriction site, generating the allelic exchange plasmid pGT-Δ*fleQ.* This plasmid was transformed into 1448A *Pseudomonas syringae* pv. phaseolicola and mutants were obtained as described in Zumaquero *et al*.^92^.

## METHOD DETAILS

### Plant growth and inoculation

*Phaseolus vulgaris* bean cultivar Canadian Wonder plants were cultivated under controlled conditions at 23°C, 95% humidity. Artificial light was maintained for periods of 16 hours within the 24 hours of the day. All experiments carried out were performed using 10-day-old plants.

For the preparation of bacterial inoculum, bacterial lawns were cultivated onto LB plates for 48 hours at 28°C. Subsequently, biomass was resuspended in 2 ml of 10 mM MgCl_2_. The optical density OD_600_ was adjusted to 0.1 corresponding to the concentration of 5 x 10^7^ colony forming units per millilitre (CFU/ml). Serial dilutions were performed to achieve the desired final inoculum concentration.

The infiltration of bean leaves for visualizing microcolonies using confocal microscopy was performed using a needleless syringe with bacterial suspension at 5·10^5^ CFU/ml. The inoculation of bean leaves for visualizing bacteria on surface using confocal microscopy was performed by dipping. For that, a bacterial suspension with 5 x 10^7^ CFU/ml was prepared in a 10 mM MgCl_2_ solution, and the entire leaf was submerged in the inoculum for a few seconds. Visualization was performed 6 hours post-inoculation (hpi).

For infiltrating bean leaves to extract bacteria from the apoplast for subsequent analysis by flow cytometry and microscopy, the method described in Rufián *et al.*^96^ was followed. This involved immersing the entire leaf in a bacterial solution with a concentration of 5 x 10^4^ CFU/ml, containing 0.01% Silwett L-77 (Crompton Europe Ltd, Evesham, UK), and using a pressure chamber. Bacteria were recovered from the plant at 4 dpi through apoplastic fluid extraction. This extraction process, as described in Rufián *et al*.^96^, entailed pressure infiltrating a whole leaf with 10 ml of a 10 mM MgCl_2_ solution inside a 20 ml syringe. After applying 5 cycles of pressure, the flow-through was collected and transferred to a fresh 50 ml tube. Three thousand microlitres of the flow-through were directly analysed by flow-cytometry. Simultaneously, the 50 ml tube were centrifuged for 30 minutes at low speed (900 *g*) at 4°C. The resulting pellets were resuspended into 1 ml of a 10 mM MgCl_2_ solution and subsequently analysed by microscopy.

To compare flagellar expression in cells mechanically extracted from or naturally exiting leaves, bean leaves were infiltrated with a 5 x 10^7^ (for 1 dpi) or 5 x 10^5^ CFU/ml (for 7 dpi) bacterial suspension using a pressure chamber, as described above. For natural exit, leaves were detached from the stem by cutting the petiole in the base of the leaf blade at the specified timepoints and incubated for 30 minutes into a 50 ml tube containing 30 ml of 10 mM MgCl_2_ to analyze natural exit. Mechanical bacterial extraction from the apoplast was carried out as described above.

### Flow Cytometry and Cell Sorting

For HIM cultures, five hundred µl of an overnight *P. syringae* LB culture was washed twice in 10 mM MgCl_2_, added to 4.5 ml of HIM and incubated at 28°C for 24h. LB cultures were obtained from an overnight incubation in LB and apoplast-extracted bacterial suspensions were obtained as indicated in Plant growth and inoculation section. Three hundred µl of the cultures in HIM, LB or in plant were analysed using a BD FACS Verse cytometer (BD Biosciences, USA) and graphs were performed with the Kaluza software (Beckman Coulter, USA). FITC-A filter was used to visualise GFP signal and PE-A filter for tdTomato signal. To ensure bleed through was not taking place, strains with transcriptional single fusions to *GFP3* and *tdTomato* were analysed with the PE-A and FITC-A filters respectively, with the observation of fluorescence as the negative control level.

For cell sorting, stationary cultures in LB obtained after an overnight incubation were sorted using a BD FACSAria^TM^ Fusion flow cytometer (BD Biosciences, USA). And exponential cultures in HIM obtained after 24 hours of incubation from 0.13 OD_600_ were sorted using a MoFloTM XDP cytometer (Beckman Coulter, USA). To initiate the process of sorting, gates were drawn to distinguish cells displaying fluorescence levels overlapping with the 1448A non-GFP bacterial population, which served as negative control, from cells expressing higher GFP levels, as indicated in the corresponding histogram. Based on this analysis, 1 x 10^5^ events were sorted for cells expressing higher GFP levels and lower GFP level. Cells from each gate were collected into separate sterile tubes. After sorting, cells were centrifuged at 12,000 *g* for 10 minutes, and the resulting pellets were resuspended into 10 mM MgCl_2_. An aliquot of sorted cells was run again at the cytometer to confirm the differences in expression between the separated populations. Data from cytometry experiments were analysed using the Kaluza Software (Beckman Coulter, USA) for further analysis and visualization.

### Confocal microscopy

For single-cell visualization of apoplast-extracted bacteria and cultured bacteria, suspensions of 2 µl were deposited over a 0.17 mm coverslip and an agar-pad square was placed on top of the drop to create a bacterial monolayer, following the method described in Rufián *et al*.^96^. To visualize all cells, bacterial suspensions were stained with FM4-64 (Life Technologies) at 20 μM, and bacterial membranes were visualized with fluorescent light, alternatively, in other cases, bright field images were included. Images of single-cell bacteria were acquired using the Zeiss LSM880 confocal microscope (Zeiss, Germany).

For the visualization of *P. syringae* microcolonies and surface cells, sections of inoculated *P. vulgaris* leaves (approximately 5 mm^2^) were carefully excised using a razor blade and mounted on slides in double-distilled H_2_O positioning the lower epidermis toward objective. A 0.17 mm coverslip was placed over the sample. Images of the leaf mesophyll and apoplast-extracted bacteria were taken using the Leica Stellaris 8 confocal microscope (Leica Microsystems GmbH, Germany) and Zeiss LSM880 confocal microscope (Zeiss, Germany).

Filters for wavelength selection were used for the visualization of the following fluorophores (excitation/ emission): eCFP (405 nm/450 to 500), GFP (488 nm/ 500 to 533 nm), FM4-64 (488 nm/ 604-674 nm), tdTomato (514 nm/570 to 600) and plant autofluorescence (514/ 605 to 670 nm). Image processing was performed using Leica LAS AF (Leica Microsystems, Germany) software. To ensure bleedthrough was not taking place, strains with transcriptional single fusions to *GFP3*, *tdTomato* and strains constitutively labelled with *eCFP* were observed under the microscope in the conditions of visualization mentioned above, with the observation of no fluorescence. Z series imaging was taken at 1 μm using 40x objectives.

### Time-lapse microscopy

Heterogeneous flagellum expression was measured during microcolony formation on HIM + 1.25% agarose pads as follows: 2X HIM medium was mixed with a melted 2.5% agarose solution and immediately placed in the wells of a custom 3D-printed mold (template available here: https://github.com/JLuneau/Pseudomonas_AgarPads_fliC/tree/main/3D_printed_AgarPad_mold) disposed on a 50 mm round coverslip (Epredia, CB005005A140MNZ0). To ensure the flatness of the pads, another coverslip was immediately placed on top of the mold. The pads were solidified for 15 minutes at room temperature. The bottom coverslip was removed and 4 µl of bacterial suspensions adjusted to OD=0.005 in HIM were dropped on the pads surface. Right after the droplets dried, a new coverslip was placed on the mold and the assembled device was mounted on the microscope. For time-lapse experiments, images were taken every 15 min, starting 4 hours after cells were placed on the pads and for 24 hours at 25°C. Images were acquired using the NIS-Elements software on a Nikon Eclipse Ti2 inverted microscope equipped with a Hamamatsu ORCA-Flash4.0LT Digital camera and a Nikon Plan Apo Lambda 100X/1.45 Oil objective. The 1.5X manual knob was engaged to enhance magnification. Illumination settings: Phase contrast, 100 ms, 50% intensity; GFP (470 nm excitation & 519 nm emission filters), 300 ms, 50% intensity. All imaging data is available upon request.

### Time-lapse image analysis

Time-lapse movies were visually inspected using Fiji 2.14.0 to crop the region of interest around microcolonies and to remove later frames when cells overlapped. Cells were segmented and tracked using the DeLTA 2.0 deep learning-based pipeline^98^ with the default pre-trained models for segmentation and tracking. Time-lapse data analysis was performed using custom Python scripts adapted from Kaczmarczyk *et al.* (2022)^99^ (available here: https://github.com/JLuneau/Pseudomonas_AgarPads_fliC). Visual inspection of DeLTA 2.0 output movies showed that while segmentation errors were rare, tracking errors were frequent at late time points. In consequence, similarly to Kaczmarczyk *et al.*^99^, we filtered out erroneous cell tracks. Upon division, *i*) we kept cells for which two sister cells were tracked for at least four frames after division, *ii*) we excluded sister cells for which the cumulated length at birth differed strongly from the length of the mother cell before division (increase or decrease of more than 20%) and *iii*) we excluded sister cells which showed unexpectedly large jumps in cell length between two frames (increase or decrease of more than 20%). For all retained cells, the *fliC* expression level was estimated as the mean fluorescent intensity in the GFP channel for all pixels belonging to a single cell, averaged over the lifetime of each individual cell. The growth rate was obtained by performing a linear regression on the log-transformed cell length over the lifetime of each cell. To estimate the cost of flagellum expression, we grouped cells into two classes: the GFP-high cells showing a mean fluorescence intensity above the median fluorescence intensity of all cells, and the GFP-low cells showing a mean fluorescence intensity below the population’s median.

### Live-dead stanning

One drop of the propidium iodide solution Ready Probes^TM^ (Thermo Fisher Scientific, USA) was added to 300 μl of the suspension with apoplast-extracted bacteria and live-dead bacteria were identified by flow-cytometry. For live-dead staining, bacteria were syringae-infiltrated with a suspension of 5 x 10^4^ CFU/ml in bean leaves and apoplast-extracted at 4 days post-inoculation.

### Competitive index (CI) assay

The competitive index (CI) assay is calculated by determining the ratio between the mutant strain and the wild type in the output sample divided by that on the input (which should be 1.0)^100–102^. Assays were performed after the mixed strains have been growing in either bean leaves or LB and HIM cultures.

Assays performed in bean plants (*Phaseolus vulgaris* cv. Canadian wonder) were carried out as detailed in Macho *et al.,* (2007)^102^. Bean plants were inoculated with 200 µl of a mixed bacterial suspension containing 5 x 10^4^ CFU/ml of each strain, consisting of an equal proportion of wild type and mutant strains. Inoculation was performed using a 1 ml syringae without needle. Samples were extracted for quantification after 4 days of post-inoculation. Bacterial recovery was carried out by taking 5 discs of 1 cm diameter from the infected leaf with a cork borer and homogenising them by mechanical disruption into 1 ml of 10 mM MgCl_2_. After homogenization, serial dilutions of the bacterial suspensions were prepared and plated onto agar plates supplemented with cycloheximide 2 μg/ml. Bacterial enumeration and CIs were calculated after 2 days of growth at 28°C. To distinguish wild type from mutant bacteria within the mixed infection, an aliquot from the same dilution was plated onto LB agar and LB agar plates supplemented with kanamycin.

For CIs assays performed in LB cultures, 500 μl of a mixed bacterial suspension with 5×10^5^ CFU/ml was inoculated into 4500 μl of LB liquid in culture tubes. For CIs assays performed in HIM cultures, 500 μl of a mixed bacterial suspension with 5×10^7^ CFU/ml was inoculated into 4500 μl of LB liquid in culture tubes. After 24 hours of incubation with continuous agitation, in both LB or HIM cultures, serial dilutions were prepared and plated onto LB agar and LB agar plates supplemented with kanamycin.

To confirm dosage and relative proportion of the strains, serial dilutions of the inoculum were plated onto LB agar and LB agar plates supplemented with the appropriate antibiotic. After bacterial counting, the ratio of the wild type *versus* the mutant strain should be close to 1. The competitive indices represent the mean of three independent experiments, each with three replicates. Error bars indicate standard error. Statistical analysis included a two-tailed Student’s t-test with a significance threshold of P < 0.05 to assess deviations from a ratio of 1.

### In vitro growth curves

Growth curves to analyse growth differences in mutant and overexpressing strains were perform in 96-well plates (Biofil, China), adjusting the bacterial inoculum to an optical density (A_600_) of 0.13 in HIM in 150 µl of final volume. The inoculum was obtained from an overnight LB culture and cells were washed twice with MgCl_2_ before adjusting the optical density. Plates were incubated for 50 hours at 28°C with agitation in a EONC plate reader (Bio Tek Instruments, USA).

Growth curves to compare Δ*hrpL* growth difference *versus* the wild type strain were performed in culture tubes in HIM with an initial optical density of 0.13 (Abs_600_). The inoculum was obtained from an overnight culture in LB, washed twice with MgCl_2_. Samples were taken at 20, 24, 26, 28, 30, 34, 38, 44, 48 and 50 hours.

To calculate bacterial growth rate, the log10 of absorbance data were calculated and represented versus time. The regression curve was calculated over the zone of exponential growth and the graph slope obtained was used as the growth rate.

### Flagellar motility assay

Flagellar motility assays conducted after the sorting of the *fliC::GFP3* strain were performed inoculating 2 µl of the aliquots obtained after the cell sorting in HIM plates containing 2.5 g/l agar or in Tryptone plates containing 3%, tryptone 5% MgCl_2_ and 2.5 g/l agar. Plates were subsequently incubated at 28°C, and digital photographs were captured to measure the diameter of the swimming halo. Measurements were calculated and the ratio of the high-expressing sorted cells to the low-expressing sorted cells was determined.

## QUANTIFICATION AND STATISTICAL ANALYSIS

All quantification and statistical analysis described in this study was performed using Prism. Details of the analysis used, and level of significance are indicated in the figure legends of each experiment.

## MULTIMEDIA TITLES

**Supplemental movie S1.** Time-lapse movie of the phase contrast of a strain carrying *hopAB1*::*GFP3* multiplying in HIM.

**Supplemental movie S2.** Time-lapse movie of the GFP fluorescence of a strain carrying *hopAB1*::*GFP3* multiplying in HIM.

**Supplemental movie S3.** Time-lapse movie of the phase contrast of a strain carrying *hrpL*::*GFP3* multiplying in HIM.

**Supplemental movie S4.** Time-lapse movie of the GFP fluorescence of a strain carrying *hrpL*::*GFP3* multiplying in HIM.

**Supplemental movie S5.** Time-lapse movie of the phase contrast of a strain carrying *fliC*::*GFP3* multiplying in HIM.

**Supplemental movie S6.** Time-lapse movie of the GFP fluorescence of a strain carrying *fliC*::*GFP3* multiplying in HIM.

**Supplemental movie S7.** Movie of the Z-stack compilation of a microcolony formed by a strain carrying *fliC*::*GFP3* in the apoplast 3 dpi.

**Supplemental movie S8.** Movie of the Z-stack compilation of a microcolony formed by a strain carrying *hopAB1*::*GFP3 fliC*::*tdT* in the apoplast 4 dpi. Individual planes are observed in Figure 5B.

**Supplemental movie S9.** Movie of the Z-stack compilation of a microcolony formed by a strain carrying *hrpL*::*GFP3 fliC*::*tdT* in the apoplast 4 dpi. Individual plane is observed in Figure 5D.

## SUPLEMENTAL INFORMATION TITLES AND LEGEND

**Figure S1. Flagella expression is heterogeneous on plant surface.**

Selected images of the *hopAB1*::*GFP3 fliC*::*tdT eCFP* strain on the plant surface at 6 hours post inoculation (hpi) by dipping the leaf into a 5 x 10^7^ CFU/ml bacterial suspension. tdTomato panel shows the fluorescence of tdTomato associated to *fliC* expression, and eCFP panel shows the fluorescence of eCFP as constitutive expression reporter, using grey scale in both cases to improve contrast. The GFP panel, corresponding to *hopAB1* expression, is not shown since no fluorescence was detected. Scale bars correspond to 20 μm. Contrast and brightness were adjusted to improve visualization but were kept constant across panels. Circles highlight bacteria detected in the eCFP panel without expression in the tdTomato channel (Flagella^OFF)^.

**Figure S2. Flagella display heterogenous expression in *Pseudomonas syringae*.**

(A) Confocal microscopic images of a strain carrying a chromosome-located *fliC*::*tdT* transcriptional fusion grown either in LB in an overnight culture (upper panels), in HIM during 24 hours (central panels), or extracted from bean leaf apoplasts 4 days post inoculation (dpi) with 5×10^4^ CFU/ml (bottom panels). tdTomato panels show the fluorescence of tdTomato as reporter of the *fliC* gene expression and BF panel corresponds to the bright field channel. Scale bars correspond to 2 µm.

(B) Flow cytometry analysis of the *fliC*::*tdT* strain obtained in the same conditions than in A. Dot plots show cell size *versus* tdTomato fluorescence intensity. Data are represented as arbitrary units in logarithmic scale. Data displayed corresponds to data collected for 100,000 events per sample. The non-tdT graph show autofluorescence levels displayed by the wild type strain not carrying any fluorescent gene marker. Vertical lines leave 99 % of the data acquired in the non-tdT strain to the left and is used as a reference to differentiate between OFF and ON cells. A and B show typical results of at least three independent replicates.

**Figure S3. T3SS and flagellar single cell expression distribution across HIM and apoplast populations.**

Dot plot graphs display the fluorescence intensity of GFP or tdTomato *versus* the cell size in the non-fluorescent bacteria (wild type strain), or the strains carrying *hrpL*::*GFP3 fliC::tdT* or *hopAB1::gfp fliC::tdT* corresponding to data shown in Figure 4 as GFP fluorescence *versus* that of tdTomato. Vertical lines leave 99 % of the data acquired for the non-fluorescent strain to the left and is used as a reference to differentiate between OFF and ON cells. Fluorescence data is represented as arbitrary units. All data was collected for 100,000 events per sample. Figure show representative results of at least three independent experiments.

(A) Bacteria grown in HIM during 24 hours after diluting an overnight grown LB culture, displaying typical bistable expression of both *hrpL*::*GFP3* and *hopAB1::gfp* and heterogeneous (occasionally bistable) expression of *fliC::tdT*. In the case of the *hopAB1::gfp fliC::tdT* strains in HIM, an additional vertical line (dashed) has been added to mark the separation between the bistable subpopulations differing in *hopAB1* expression, which is higher that the line established using the non-fluorescent strain on the account of the high basal expression levels of this gene.

(B) Bacteria extracted from bean leaf apoplasts 4 days post inoculation (dpi) with 5 x 10^4^ CFU/ml displaying typical heterogeneous (never bistable) expression of *hrpL*::*GFP3, hopAB1::gfp* and *fliC::tdT*.

**Figure S4. T3SS and flagella expression impact on bacterial growth and growth cost associated with single cell levels of flagella production is not due to GFP accumulation.**

(A) Growth rates of the 1448A wild type strain, a derivative carrying: a plasmid that determines constitutive expression of HrpL (pHrpL), or of fleQ (pFleQ), or a deletion of the fleQ gene (ΔfleQ), in HIM.

(B) Selected timelapse images of the constitutively expressing GFP (eGFP) strain during the microcolony development on agar pads in phase contrast (top) and GFP fluorescence (bottom) channels. Contrast and brightness were adjusted to improve visualization but were kept constant across frames of each timelapse.

(C) Comparisons of mean growth rate for the *eGFP* strain and for the *fliC*::*GFP3* strain. Note that GFP in the *eGFP* strain has a much higher fluorescence intensity compared to *fliC*::*GFP3*. *eGFP* shows a significantly lower average growth rate compared to *fliC*::*GFP* (Mann-Whitney U test, *P* = 7.7 x 10^-15^).

(D) Comparison of growth rate of cells with high and low GFP signal (determined by splitting the population into two groups using the median fluorescence intensity value) for the eGFP strain shows no significant differences between the growth rates of cells cells expressing high or low GFP levels (Mann-Whitney U test, P<0.05).

(E) Correlation between growth rates and fluorescence intensity of individual fliC::GFP3 cells indicates that higher fliC expression is associated with slower growth. The shaded area shows the 95% confidence interval.

(F) Correlation between growth rates and fluorescence intensity of individual eGFP cells indicates that higher GFP expression is not associated with slower growth. The shaded area shows the 95% confidence interval.

**Figure S5. Dead-live staining shows neither bias towards Flagella^ON^ cells nor for T3SS^OFF^ cells during plant growth.**

Graph shows the dead/live ratio for bacteria expressing either expressing or not *fliC*::*GFP3, hrpL*::*GFP3*, *hopAB1*:*GFP3* or *hrcU*::*GFP3* during plant growth. Apoplast-extracted bacteria at 4 days post-inoculation in bean leaves were stained with a solution of Propidium iodide (PI). Data were obtained by flow cytometry analysis and GFP levels were used to differentiate between ON and OFF cells using the non-fluorescent wild type strain as reference, as indicated before. Fluorescence of PI was used to differentiate dead and alive bacteria comparing to the non-fluorescent wild type. Each dot represents an extraction event and therefore a different biological replicate.

## Notes

### Competing Interest Statement

The authors have declared no competing interest.

