## Supplemental material for "Cooperative colonization of the host and pathogen dissemination involves stochastic and spatially structured expression of virulence traits"

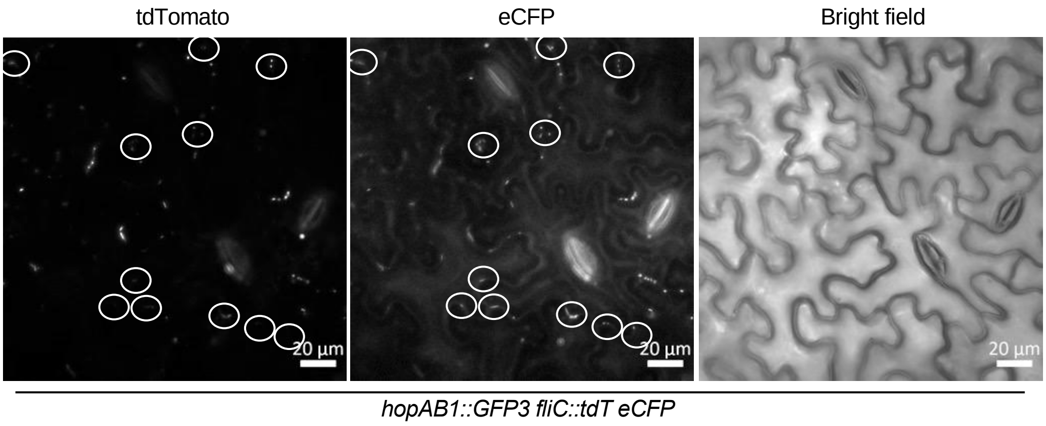


**Figure S1. Flagella expression is heterogeneous on plant surface.**

Selected images of the *hopAB1*::*GFP3 fliC*::*tdT eCFP* strain on the plant surface at 6 hours post inoculation (hpi) by dipping the leaf into a 5 x $10^7$ CFU/ml bacterial suspension. tdTomato panel shows the fluorescence of tdTomato associated to *fliC* expression, and eCFP panel shows the fluorescence of eCFP as constitutive expression reporter, using grey scale in both cases to improve contrast. The GFP panel, corresponding to *hopAB1* expression, is not shown since no fluorescence was detected. Scale bars correspond to 20 µm. Contrast and brightness were adjusted to improve visualization but were kept constant across panels. Circles highlight bacteria detected in the eCFP panel without expression in the tdTomato channel (Flagella$^{OFF)}$.

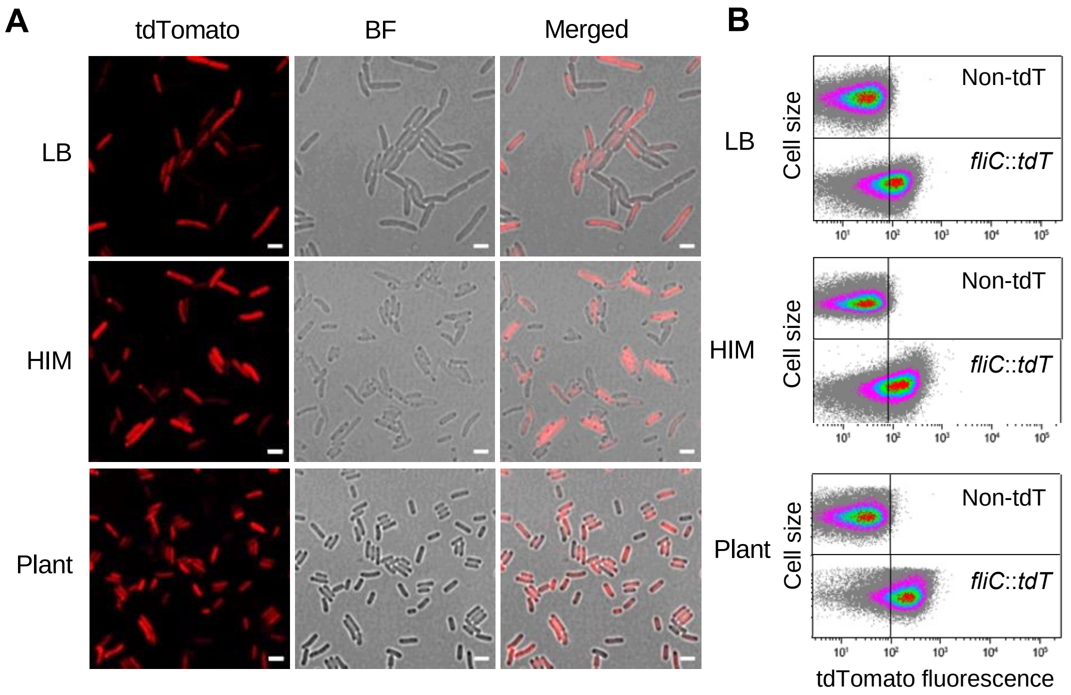


**Figure S2. Flagella display heterogenous expression in *Pseudomonas syringae*.**

(A) Confocal microscopic images of a strain carrying a chromosome-located *fliC*::*tdT* transcriptional fusion grown either in LB in an overnight culture (upper panels), in HIM during 24 hours (central panels), or extracted from bean leaf apoplasts 4 days post inoculation (dpi) with $5x10^4$ CFU/ml (bottom panels). tdTomato panels show the fluorescence of tdTomato as reporter of the *fliC* gene expression and BF panel corresponds to the bright field channel. Scale bars correspond to 2 µm.

(B) Flow cytometry analysis of the *fliC*::*tdT* strain obtained in the same conditions than in A. Dot plots show cell size *versus* tdTomato fluorescence intensity. Data are represented as arbitrary units in logarithmic scale. Data displayed corresponds to data collected for 100,000 events per sample. The non-tdT graph show autofluorescence levels displayed by the wild type strain not carrying any fluorescent gene marker. Vertical lines leave 99 % of the data acquired in the non-tdT strain to the left and is used as a reference to differentiate between OFF and ON cells. A and B show typical results of at least three independent replicates.

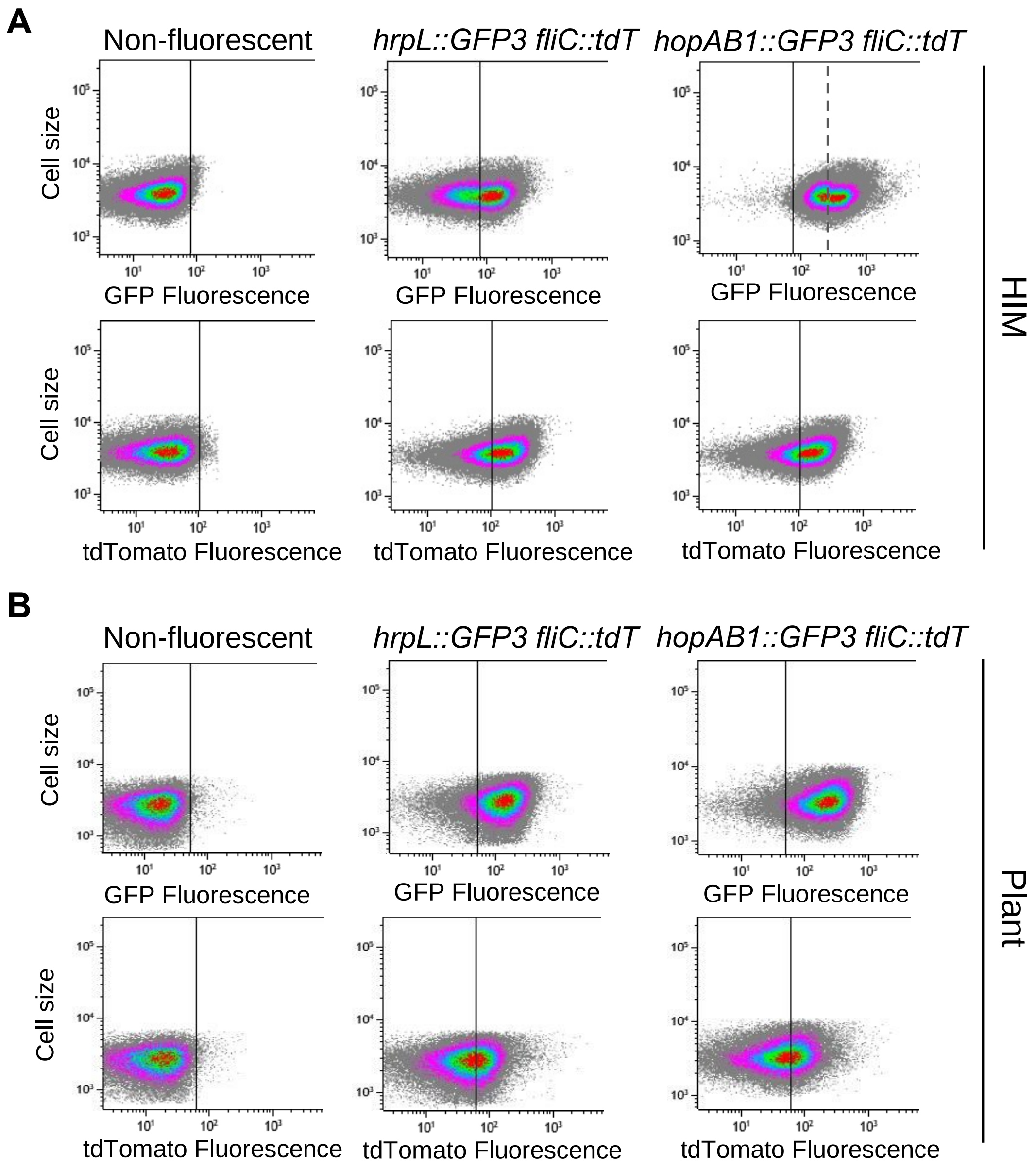


**Figure S3. T3SS and flagellar single cell expression distribution across HIM and apoplast populations.**

Dot plot graphs display the fluorescence intensity of GFP or tdTomato *versus* the cell size in the non-fluorescent bacteria (wild type strain), or the strains carrying *hrpL*::*GFP3 fliC::tdT* or *hopAB1::gfp fliC::tdT* corresponding to data shown in Figure 4 as GFP fluorescence *versus* that of tdTomato. Vertical lines leave 99 % of the data acquired for the non-fluorescent strain to the left and is used as a reference to differentiate between OFF and ON cells. Fluorescence data is represented as arbitrary units. All data was collected for 100,000 events per sample. Figure show representative results of at least three independent experiments.

(A) Bacteria grown in HIM during 24 hours after diluting an overnight grown LB culture, displaying typical bistable expression of both *hrpL*::*GFP3* and *hopAB1::gfp* and heterogeneous (occasionally bistable) expression of *fliC::tdT*. In the case of the *hopAB1::gfp fliC::tdT* strains in HIM, an additional vertical line (dashed) has been added to mark the separation between the bistable subpopulations differing in *hopAB1* expression, which is higher that the line established using the non-fluorescent strain on the account of the high basal expression levels of this gene.
(B) Bacteria extracted from bean leaf apoplasts 4 days post inoculation (dpi) with 5 x $10^4$ CFU/ml displaying typical heterogeneous (never bistable) expression of *hrpL*::*GFP3, hopAB1::gfp* and *fliC::tdT*.

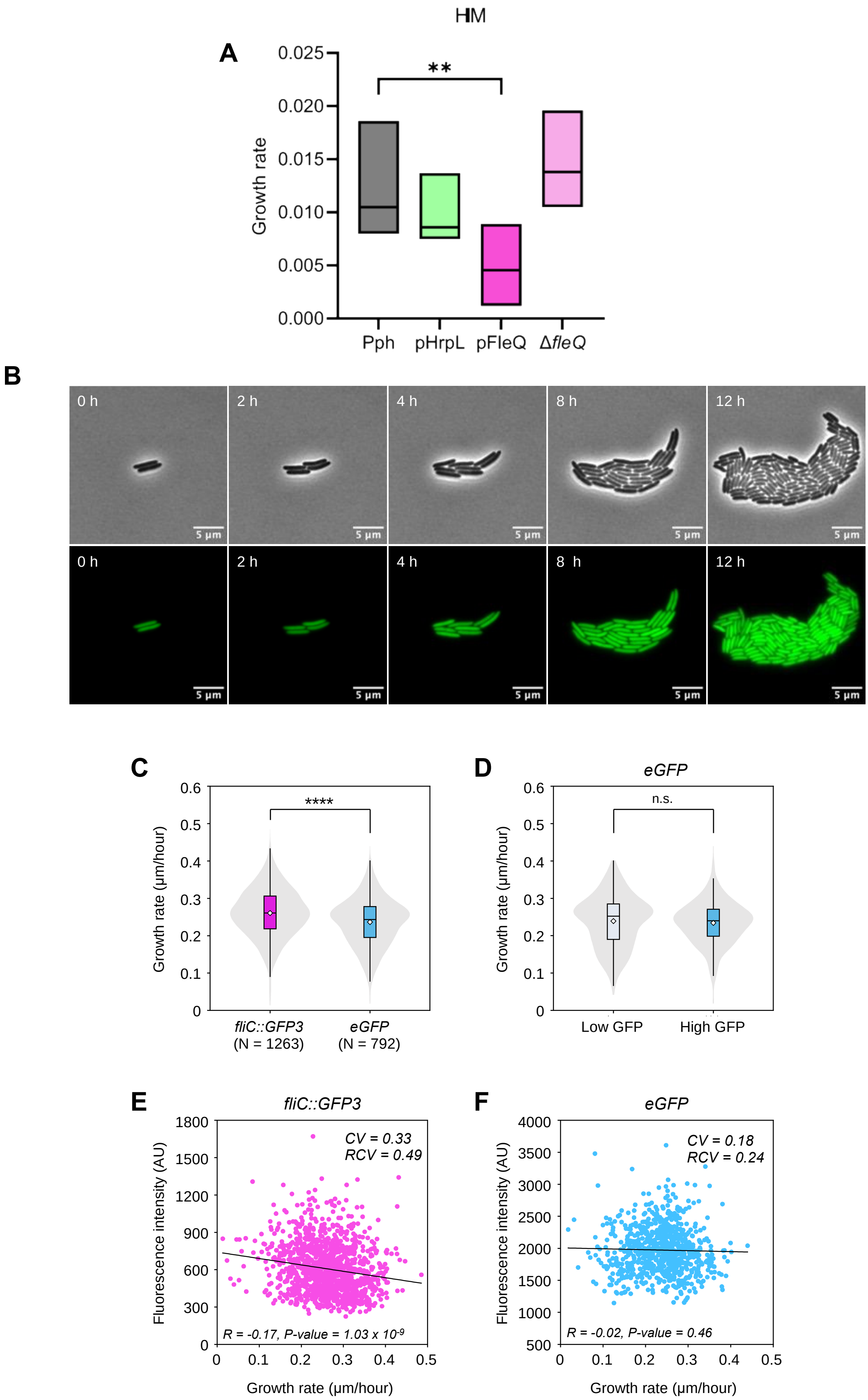
HM
A
0.025
0.020
0.015
0.010
0.005
0.000
Growth rate
**
Pph
pHrpL
pFleQ
Δ*fleQ*
B
0 h
2 h
4 h
8 h
12 h
5 µm
C
0.6
0.5
0.4
0.3
0.2
0.1
0
Growth rate (µm/hour)
****
*fliC::GFP3*
(N = 1263)
*eGFP*
(N = 792)
D
*eGFP*
n.s.
Growth rate (µm/hour)
Low GFP
High GFP
E
*fliC::GFP3*
1800
1500
1200
900
600
300
0
Fluorescence intensity (AU)
*CV = 0.33*
*RCV = 0.49*
*R = -0.17, P-value = 1.03 x 10-9*
0
0.1
0.2
0.3
0.4
0.5
Growth rate (µm/hour)
F
*eGFP*
4000
3500
3000
2500
2000
1500
1000
500
Fluorescence intensity (AU)
*CV = 0.18*
*RCV = 0.24*
*R = -0.02, P-value = 0.46*
Growth rate (µm/hour)

**Figure S4. T3SS and flagella expression impact on bacterial growth and growth cost associated with single cell levels of flagella production is not due to GFP accumulation.**

(A) Growth rates of the 1448A wild type strain, a derivative carrying: a plasmid that determines constitutive expression of HrpL (pHrpL), or of fleQ (pFleQ), or a deletion of the fleQ gene (ΔfleQ), in HIM.

(B) Selected timelapse images of the constitutively expressing GFP (eGFP) strain during the microcolony development on agar pads in phase contrast (top) and GFP fluorescence (bottom) channels. Contrast and brightness were adjusted to improve visualization but were kept constant across frames of each timelapse.

(C) Comparisons of mean growth rate for the *eGFP* strain and for the *fliC*::*GFP3* strain. Note that GFP in the *eGFP* strain has a much higher fluorescence intensity compared to *fliC*::*GFP3*. *eGFP* shows a significantly lower average growth rate compared to *fliC*::*GFP* (Mann-Whitney U test, $P = 7.7 \times 10^{-15}$).

(D) Comparison of growth rate of cells with high and low GFP signal (determined by splitting the population into two groups using the median fluorescence intensity value) for the eGFP strain shows no significant differences between the growth rates of cells cells expressing high or low GFP levels (Mann-Whitney U test, $P<0.05$).

(E) Correlation between growth rates and fluorescence intensity of individual fliC::GFP3 cells indicates that higher fliC expression is associated with slower growth. The shaded area shows the 95% confidence interval.

(F) Correlation between growth rates and fluorescence intensity of individual eGFP cells indicates that higher GFP expression is not associated with slower growth. The shaded area shows the 95% confidence interval.

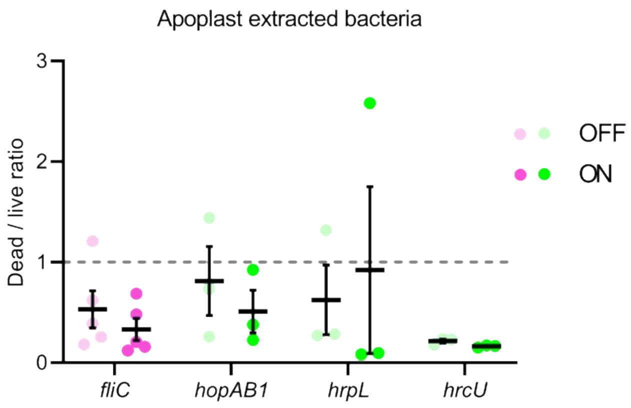


**Figure S5. Dead-live staining shows neither bias towards Flagella$^{ON}$ cells nor for T3SS$^{OFF}$ cells during plant growth.**

Graph shows the dead/live ratio for bacteria expressing either expressing or not *fliC*::*GFP3, hrpL*::*GFP3*, *hopAB1*:*GFP3* or *hrcU*::*GFP3* during plant growth. Apoplast-extracted bacteria at 4 days post-inoculation in bean leaves were stained with a solution of Propidium iodide (PI). Data were obtained by flow cytometry analysis and GFP levels were used to differentiate between ON and OFF cells using the non-fluorescent wild type strain as reference, as indicated before. Fluorescence of PI was used to differentiate dead and alive bacteria comparing to the non-fluorescent wild type. Each dot represents an extraction event and therefore a different biological replicate.
